# Hearing function moderates age-related changes in brain morphometry in the HCP Aging cohort

**DOI:** 10.1101/2024.04.22.590589

**Authors:** Robert M. Kirschen, Amber M. Leaver

**Affiliations:** Department of Radiology, Northwestern University, Chicago, IL, 60611

**Author notes:** Corresponding Author: Amber M. Leaver Ph.D. Address: 737 N Michigan Ave Suite 1600 Chicago, IL 60611.

**Keywords:** Hearing Loss, Aging, Temporal Cortex, Morphometry

## Abstract

**Introduction:** There are well-established relationships between aging and neurodegenerative changes, and between aging and hearing loss. The goal of this study was to determine how structural brain aging is influenced by hearing loss.

**Methods:** Human Connectome Project Aging (HCP-A) data were analyzed, including T1-weighted MRI and Words in Noise (WIN) thresholds (n=623). Freesurfer extracted gray and white matter volume, and cortical thickness, area, and curvature. Linear regression models targeted (1) interactions between age and WIN threshold and (2) correlations with WIN threshold adjusted for age, both corrected for false discovery rate (pFDR<0.05).

**Results:** WIN threshold moderated age-related increase in volume in bilateral inferior lateral ventricles, with higher threshold associated with increased age-related ventricle expansion. Age-related deterioration in occipital cortex was also increased with higher WIN thresholds. When controlling for age, high WIN threshold was correlated with reduced cortical thickness in Heschl’s gyrus, calcarine sulcus, and other sensory regions, and reduced temporal lobe white matter. Older volunteers with poorer hearing and cognitive scores had the lowest volume in left parahippocampal white matter.

**Conclusions:** Preserved hearing abilities in aging associated with a reduction of age-related changes to medial temporal lobe, and preserved hearing at any age associated with preserved cortical tissue in auditory and other sensory regions. Future longitudinal studies are needed to assess the causal nature of these relationships, but these results indicate interventions which preserve hearing function may combat some neurodegenerative changes in aging.

**KEY POINTS (3 summarizing key messages):**

1. Poorer hearing was associated with increased age-related ventricle expansion in medial temporal lobes, and reduced temporal lobe white matter at any age.
2. Poorer hearing was associated with thinner cortex in Heschl’s gyrus thickness, calcarine sulcus, and other sensory regions.
3. Preserving hearing may reduce brain aging in medial temporal lobe.

## INTRODUCTION

There are established hallmarks of brain aging on the macro-scale, including cortical and subcortical atrophy, increased ventricle size, changes in cerebral perfusion, and other specific markers associated with age-related neurodegenerative disease [Cole, 2020; Fjell et al., 2013; Frangou et al., 2022; Juttukonda et al., 2021]. Though these overall patterns of change are well characterized, there is sufficient variability to suggest that not all people experience these changes at the same rate or to the same degree [Cox and Deary, 2022]. Understanding why some people’s brains age more or faster than others could help identify risk factors and interventions to promote healthy brain aging.

Hearing loss is also a well-established hallmark of aging. Age-related hearing loss is associated with stiffening of outer hair cells in the cochlea, cumulative otologic injury from loud sounds or drugs, and other factors causing loss of peripheral input from the inner ear to the central auditory system. Hearing loss is associated with social, occupational, cognitive, and mental health impacts, including increased risk of dementia [Killeen et al., 2023; Lin et al., 2013; Stevenson et al., 2022] and depression [Li et al., 2014; Parravano et al., 2021]. Yet, hearing loss also occurs in younger adults (with risk increasing in recent years [Dillard et al., 2022]), and may impact the brain independently of age in some brain systems, while accelerating the pace of age-related changes in others. Therefore, studying the impact of hearing on the brain and on brain aging appears critical.

Hearing loss has been linked to structural changes in the central auditory system, including reduced gray and white matter in the temporal lobe [Armstrong et al., 2019; Eckert et al., 2012; Eckert et al., 2019; Li et al., 2023], including Heschl’s gyrus, the location of core/primary auditory cortex; [Eckert et al., 2019; Lin et al., 2014; Peelle et al., 2011]. Changes in other brain systems have also been identified, sometimes interpreted as compensatory changes (e.g., frontal cortex [Husain et al., 2011; Khan et al., 2021; Koops et al., 2020; Melcher et al., 2013; Qian et al., 2017; Yang et al., 2014]). However, effects of age are not always statistically dissociated from effects of hearing loss in these studies, which may also have limited sample size and age ranges.

The Human Connectome Project (HCP) offers a unique opportunity to address this issue on a much larger scale, with high-quality, well-characterized multimodal MRI datasets collected in volunteers across the lifespan [Elam et al., 2021; Harms et al., 2018]. Critically, these datasets also include a basic hearing test as part of an extended cognitive/perceptual test battery [Barch et al., 2013]. This Words in Noise (WIN) task assesses speech reception threshold, where volunteers identify spoken words embedded in multi-speaker babble. In typical populations, thresholds on WIN tasks are correlated with pure-tone thresholds assessed via air conduction, though the strength of this correlation depends on the population studied and the manner of WIN task administration [Fitzgerald et al., 2023; Holmes and Griffiths, 2019; Humes, 2021; Kam and Fu, 2020; Leaver, 2024; Vermiglio et al., 2020]. Although difficulty hearing in noisy environments can sometimes persist after hearing aid use [Davidson et al., 2022] and may indicate central hearing loss [C. Kohrman et al., 2020], these phenomena appear to be occur less frequently than peripheral/sensorineural hearing loss [Spankovich et al., 2018]. Therefore, in databases like the HCP that do not select for or exclude hearing loss during recruitment, WIN threshold is most likely to reflect peripheral hearing loss. Under this premise, the current study used WIN threshold as a proxy for peripheral hearing loss to measure the effects of hearing function on brain aging.

In the present study, we analyzed relationships between brain structure, hearing loss, and age using the HCP Aging dataset. Our goal was to identify instances where hearing function moderated the effects of aging on brain structure (i.e., a WIN-by-age interaction on brain structure). In addition, we hypothesized that hearing loss could affect the brain at any age, and therefore also identified instances where hearing loss correlated with brain structure while controlling for age (i.e., main effect of WIN threshold). Brain structure was measured comprehensively using Freesurfer pipelines on T1-weighted MRI scans, and included volume of ventricles, subcortical structures, and cortical gray and white matter, as well as cortical thickness, curvature, and surface area. Hearing loss was assessed using WIN task threshold, and exploratory analyses used the Montreal Cognitive Assessment to address the influence of cognitive status on relationships between hearing function, age and brain morphometry.

## MATERIALS & METHODS

### Participants and Data

Data for this analysis were taken from the Human Connectome Project (HCP) Aging dataset [Bookheimer et al., 2019]. Data were downloaded in August 2023 from the NIMH Data Archive, and reflect data release 2.0. In the HCP Aging study, participants underwent an MRI protocol, a NIH Toolbox battery including the Words in Noise (WIN) task, and other assessments at four sites: Washington University St. Louis, University of Minnesota, Massachusetts General Hospital, and University of California, Los Angeles. Ages ranged from 36 years to 90+. In this download, 725 participants had structural MRI data and Montreal Cognitive Assessment (MoCA) scores [Nasreddine et al., 2005], while 633 had NIH Toolbox WIN data. For the current analysis, we retained complete cases (i.e., data from participants with both MRI and WIN data) with age less than 90. Participants over 90 (n=13) were coded as the same age in the HCP dataset (1200 months), and thus were excluded from this analysis due to potential influence of inaccurate age data on statistical model fit. This yielded 622 complete cases for analysis.

### NIH Toolbox Words In Noise Task

During the Words in Noise (WIN) task, volunteers were asked to repeat words presented unilaterally (i.e., separately to each ear), spoken by one target speaker along with multi-speaker babble background noise [Zecker et al., 2013]. Signal-to-noise ratio (SNR) of target speaker to noise was varied (24, 20, 16, 12, 8, 4, 0 dB SNR), and 6 trials were presented at each SNR. In this task, the experimenter records spoken responses from the volunteer using a tablet device, and sounds are played through over-ear headphones (e.g., Sennheiser 280 Pro) with tablet volume set at a “comfortable level.” For MRI analyses, we used WIN threshold averaged over both ear as reported by NIH Toolbox.

### MRI Acquisition & Preprocessing

MRI imaging was performed at all four sites using the same hardware, a Siemens 3T Prisma [Harms et al., 2018]. The current study analyzed T1-weighted structural scans, though diffusion, perfusion, and functional MRI data are also available from the HCP-A dataset [Bookheimer et al., 2019]. Freesurfer’s reconall pipeline (Version 7.20.0, [Dale et al., 1999]) was used to extract gray and white matter volume, as well as cortical thickness, area, and curvature using T1-weighted MRI scans [Destrieux et al., 2010; Fischl et al., 2002; Fischl and Dale, 2000]. Generally speaking, metrics that are expected to decrease with age include gray and white matter volume, cortical thickness, and (perhaps to a lesser extent, [Winkler et al., 2018]) cortical surface area, while ventricle volume and mean cortical curvature are expected to increase with age, though some regions may deviate from this general pattern [Salat et al., 2004]. Each metric type (gray matter volume, white matter volume, cortical thickness, cortical area, and cortical curvature) was harmonized across study sites applying neuroCombat separately for each metric type [Fortin et al., 2017] in R (https://www.r-project.org). Extreme outliers greater than 4 standard deviations above or below the sample mean were excluded from analysis.

### Statistical Analyses

All statistical analyses were completed in R (https://www.r-project.org). To test for relationships amongst WIN threshold, site, demographic, and other variables, Pearson’s correlation, Student’s t-test, ANOVA, or Chi-squared tests were used as appropriate. For MRI analyses, linear regression models measured relationships between brain morphometry, WIN thresholds, and age (all linear factors) adjusted for participant sex (categorical factor). Two statistical models were applied. The first model was a moderation analysis targeting an interaction between age and WIN threshold, with the goal of identifying instances where age-related differences in brain morphometry differed across WIN thresholds (i.e., how brain aging is impacted by hearing function). A second, separate model targeted main effects of WIN threshold while controlling for age and sex (i.e., how hearing function impacts brain structure independent of age). For both models, effect size is reported as partial r^2^ (partial_r2 function, [Cinelli and Hazlett, 2020]), an estimate of the unique variance in freesurfer metric explained by each model term (i.e., WIN, age, or interaction). Two statistical thresholds were used: false discovery rate q< 0.05 across all metrics, and uncorrected p < 0.05 for auditory cortical regions. For main effects meeting these statistical criteria for model two, interaction effects from the moderation model are also reported. For moderation or main effects pFDR < 0.05, an exploratory analysis tested for a triple interaction between WIN threshold, age, and Montreal Cognitive Assessment (MoCA) score adjusted for sex, puncorr < 0.05

## RESULTS

### Participant characteristics and WIN threshold

Age and sex did not differ across sites (F(3,619)=2.34, p=0.07 and !^2^(3)=2.79, p=0.42, respectively), though the UCLA cohort was slightly younger (**Table 1**). Reported WIN threshold differed across sites (F(3,619)=8.83, p=0.00005), with the Minnesota cohort having slightly higher thresholds (*pTukeyHSD* <0.001 for all). As expected, WIN threshold correlated with age (r=0.55, p=0.37x10^-51^) (**Figure 1**). WIN threshold was also slightly higher on average in males (t(621)=4.03, p=0.00006; mean difference (95% CI) = 1.40(0.68) dB SNR) and for left ear stimuli (t(622)=2.66,p=0.008; mean difference (95% CI) = 0.33(0.24) dB SNR).

**Figure 1.**
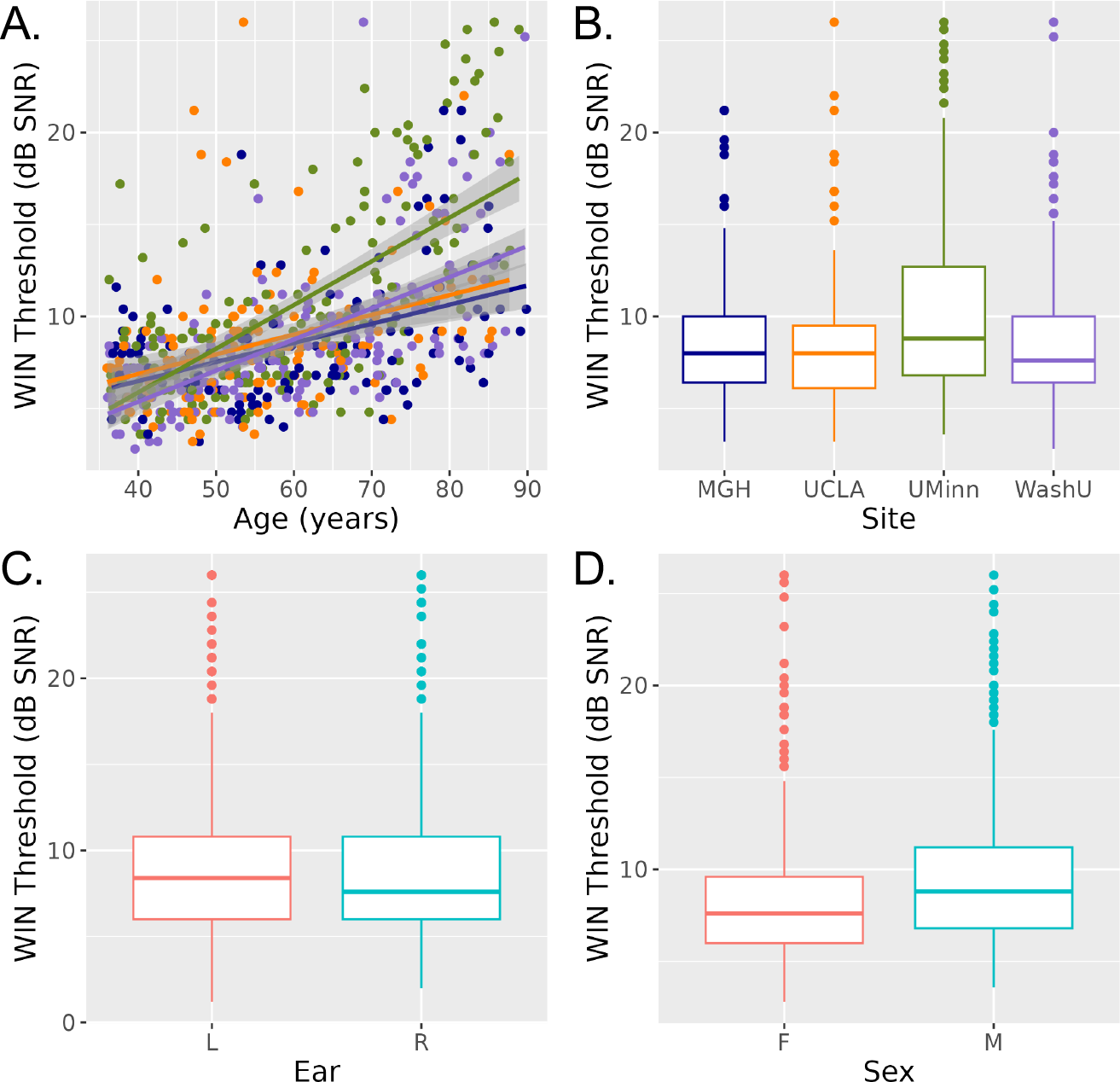
Words In Noise (WIN) Threshold differs over age, study site, ear, and sex. **A.** Scatter plot displays WIN Threshold and age for each volunteer, with color reflecting study site (MGH blue, UCLA orange, UMinn green, WashU purple). Linear regression lines are fitted for each site, with shading reflecting standard error. **B-D.** Boxplots display WIN Threshold across sites, ear, and sex at birth, respectively. Abbreviations: L, left; R, right; F, female; M, male; dB, decibel; SNR, signal to noise ratio.

**Table 1.** Demographic and Clinical Information by Site.

|  | MGH | UCLA | UMinn | WashU |
| --- | --- | --- | --- | --- |
| Sample size | 125 | 130 | 186 | 182 |
| Age, mean (SD) yrs | 60.44 (15.47) | 56.08 (12.76) | 59.62 (14.76) | 59.34 (14.47) |
| Sex, females/males | 64/61 | 78/52 | 106/80 | 109/73 |
| WIN Threshold Right Ear, mean (SD) dB SNR | 8.62 (3.75) | 8.59 (4.45) | 10.31 (5.53) | 8.29 (3.98) |
| WIN Threshold Left Ear, mean (SD) dB SNR | 8.6 (3.99) | 8.6 (3.69) | 10.76 (5.4) | 8.98 (4.37) |

### Hearing function moderates effects of age on brain structure

In MRI analyses of the effects of hearing loss on age-related differences in brain structure, interactions between WIN threshold and age were noted bilaterally in inferior lateral ventricles, such that the rate of age-related increase in volume was greater in people with higher WIN thresholds (**Figure 2A**, **Table 2**). WIN-by-age interactions were also present in occipital cortex, where the rate of age-related thinning was greater in people with higher WIN thresholds (**Figure 2B**, **Table 2**). This included several left lateral occipital regions, left cuneus, and bilateral occipital pole. In anterior cingulate cortex thickness, an interaction showed the opposite pattern: age-related decreases were more pronounced in people with better WIN thresholds.

**Figure 2.**
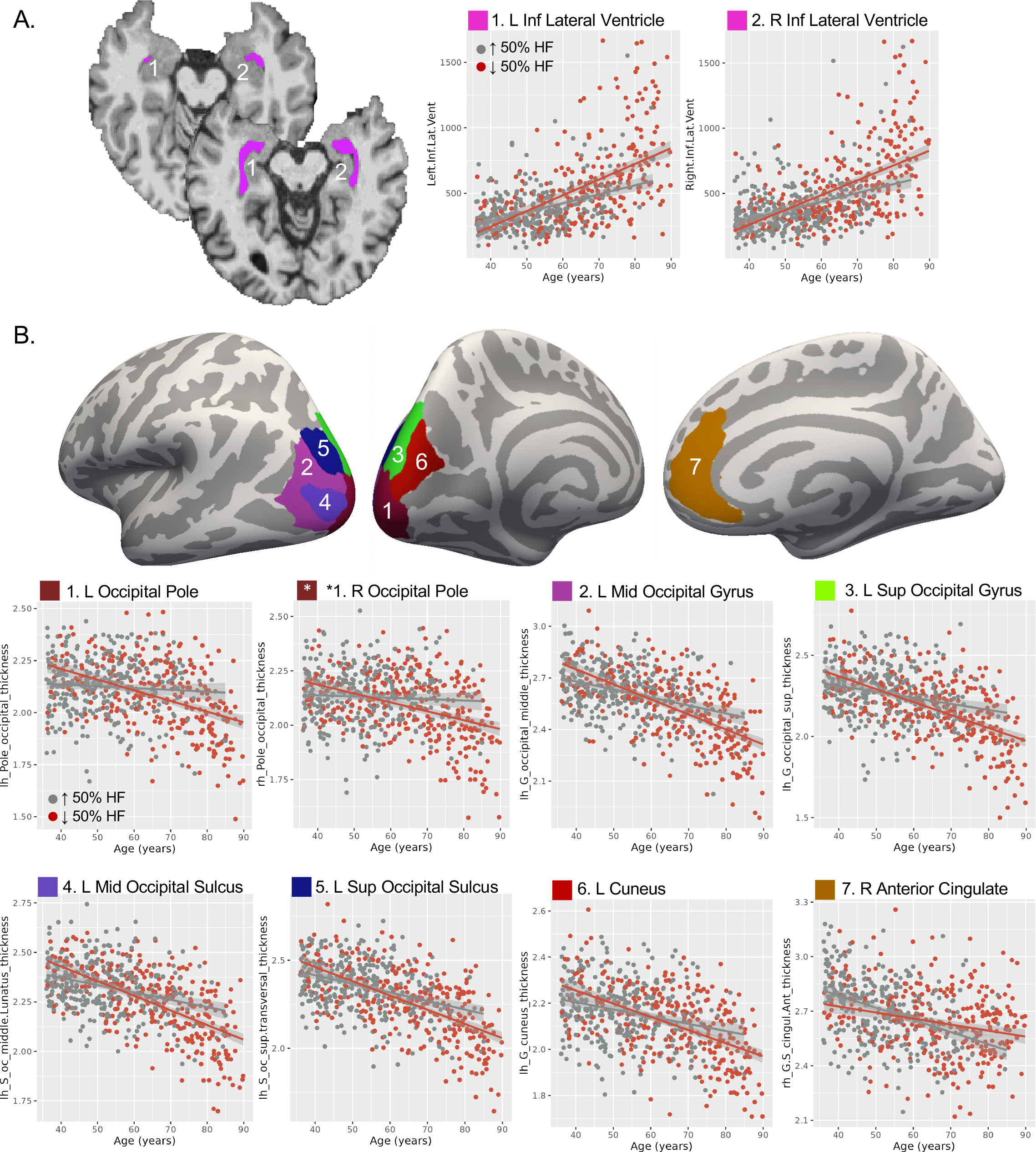
WIN Threshold moderates age-related effects on brain structure. **A.** Left and right inferior lateral ventricles exhibited a WIN-by-age interaction. Location is displayed in two representative volunteers with typical (top) and enlarged (bottom) ventricles for reference. Scatter plots at right show ventricle volume (mm^3^) and age for each volunteer, with linear regression fit displayed separately for the highest 50% of WIN thresholds (hearing loss, red) and lowest 50% of WIN thresholds (better hearing, gray). Shading reflects standard error. Note that WIN threshold was binarized for display purposes only; statistics used full range of WIN thresholds. **B.** Cortical thickness in occipital regions and right anterior cingulate cortex also showed WIN-by-age interactions, and are displayed on a template cortical surface (fsaverage). Scatter plots display cortical thickness (mm) and age for each volunteer, with regression lines plotted as in A. Abbreviations: L, left; R, right; Inf, inferior; Mid, middle; Sup, superior.

**Table 2.** WIN-by-Age Interactions on Brain Structure, p-fdr < 0.05.

| Region, Measure | effect | $\beta$ | $\beta$ SE | t | df | p | pFDR | partial $r^2$ |
| --- | --- | --- | --- | --- | --- | --- | --- | --- |
| L Inferior Lateral Ventricle, Volume | WIN*Age | 0.043 | 0.012 | 3.711 | 611 | 0.0002 | 0.02 | 0.022 |
|  | WIN | -30.868 | 9.809 | -3.147 | 611 | 0.002 | 0.24 | 0.016 |
|  | Age | 0.428 | 0.117 | 3.654 | 611 | 0.0003 | 0.000000 | 0.021 |
| R Inferior Lateral Ventricle, Volume | WIN*Age | 0.043 | 0.012 | 3.626 | 614 | 0.0003 | 0.02 | 0.021 |
|  | WIN | -29.607 | 10.094 | -2.933 | 614 | 0.003 | 0.12 | 0.014 |
|  | Age | 0.376 | 0.120 | 3.123 | 614 | 0.0019 | 0.000000 | 0.016 |
| L Cuneus Gyrus, Thickness | WIN*Age | -0.00003 | 0.00001 | -4.368 | 618 | 0.00001 | 0.002 | 0.030 |
|  | WIN | 0.023 | 0.006 | 3.724 | 618 | 0.0002 | 0.18 | 0.022 |
|  | Age | -0.00006 | 0.00007 | -0.874 | 618 | 0.38 | 0.000000 | 0.001 |
| L Middle Occipital Gyrus, Thickness | WIN*Age | -0.00004 | 0.00001 | -5.204 | 618 | 0.000000 | 0.00007 | 0.042 |
|  | WIN | 0.031 | 0.007 | 4.379 | 618 | 0.00001 | 0.09 | 0.030 |
|  | Age | -0.00015 | 0.00008 | -1.785 | 618 | 0.07 | 0.000000 | 0.005 |
| L Superior Occipital Gyrus, Thickness | WIN*Age | -0.00004 | 0.00001 | -4.010 | 618 | 0.00007 | 0.006 | 0.025 |
|  | WIN | 0.02523 | 0.00778 | 3.244 | 618 | 0.001 | 0.09 | 0.017 |
|  | Age | -0.0002 | 0.00009 | -1.635 | 618 | 0.10 | 0.000000 | 0.004 |
| L Occipital Pole, Thickness | WIN*Age | -0.00004 | 0.00001 | -5.644 | 618 | 0.000000 | 0.00001 | 0.049 |
|  | WIN | 0.033 | 0.007 | 4.944 | 618 | 0.000001 | 0.17 | 0.038 |
|  | Age | 0.0002 | 0.00008 | 1.971 | 618 | 0.05 | 0.000000 | 0.006 |
| R Occipital Pole, Thickness | WIN*Age | -0.00003 | 0.00001 | -4.067 | 618 | 0.00005 | 0.01 | 0.026 |
|  | WIN | 0.021 | 0.006 | 3.274 | 618 | 0.001 | 0.09 | 0.017 |
|  | Age | 0.00009 | 0.00008 | 1.177 | 618 | 0.24 | 0.000005 | 0.002 |
| L Middle Occipital Sulcus and Lunatus, Thickness | WIN*Age | -0.00003 | 0.00001 | -4.587 | 617 | 0.00001 | 0.001 | 0.033 |
|  | WIN | 0.024 | 0.006 | 3.807 | 617 | 0.0002 | 0.09 | 0.023 |
|  | Age | -0.00016 | 0.00007 | -2.119 | 617 | 0.03 | 0.000000 | 0.007 |
| L Sup Occipital Sulcus and Transversal, Thickness | WIN*Age | -0.00003 | 0.00001 | -3.921 | 617 | 0.0001 | 0.01 | 0.024 |
|  | WIN | 0.022 | 0.007 | 3.199 | 617 | 0.001 | 0.11 | 0.016 |
|  | Age | -0.00024 | 0.00008 | -2.830 | 617 | 0.005 | 0.000000 | 0.013 |
| R Anterior Cingulate Gyrus and Sulcus, Thickness | WIN*Age | 0.00003 | 0.00001 | 3.350 | 618 | 0.0009 | 0.05 | 0.018 |
|  | WIN | -0.023 | 0.008 | -2.813 | 618 | 0.005 | 0.25 | 0.013 |
|  | Age | -0.001 | 0.000 | -7.412 | 618 | 0.000000 | 0.000000 | 0.082 |

Note: Values listed as 0.000000 are less than 0.000001.

### Effects of hearing loss on brain structure controlling for age

Main effects of WIN threshold independent of age were noted in many temporal lobe regions (**Figure 3**, **Table 3**). This included cortical thickness in left Heschl’s gyrus, where high WIN threshold was associated with less tissue. Higher WIN threshold was similarly associated with less volume in left entorhinal and parahippocampal white matter, as well as right middle temporal and fusiform white matter. Calcarine sulcus thickness also showed a similar pattern, though a significant WIN-by-age interaction was also noted in this metric (t(618)=-3.07, p=0.002, partial r^2^=0.015). In right rectus gyrus (medial orbitofrontal cortex) thickness and left mid-posterior cingulate cortex curvature, positive correlations were present, where poorer performance associated with higher morphometry measures. Of all these structures, only calcarine sulcus thickness showed a WIN-by-age interaction puncorr < 0.05 (**Supplementary Table 1**).

**Figure 3.**
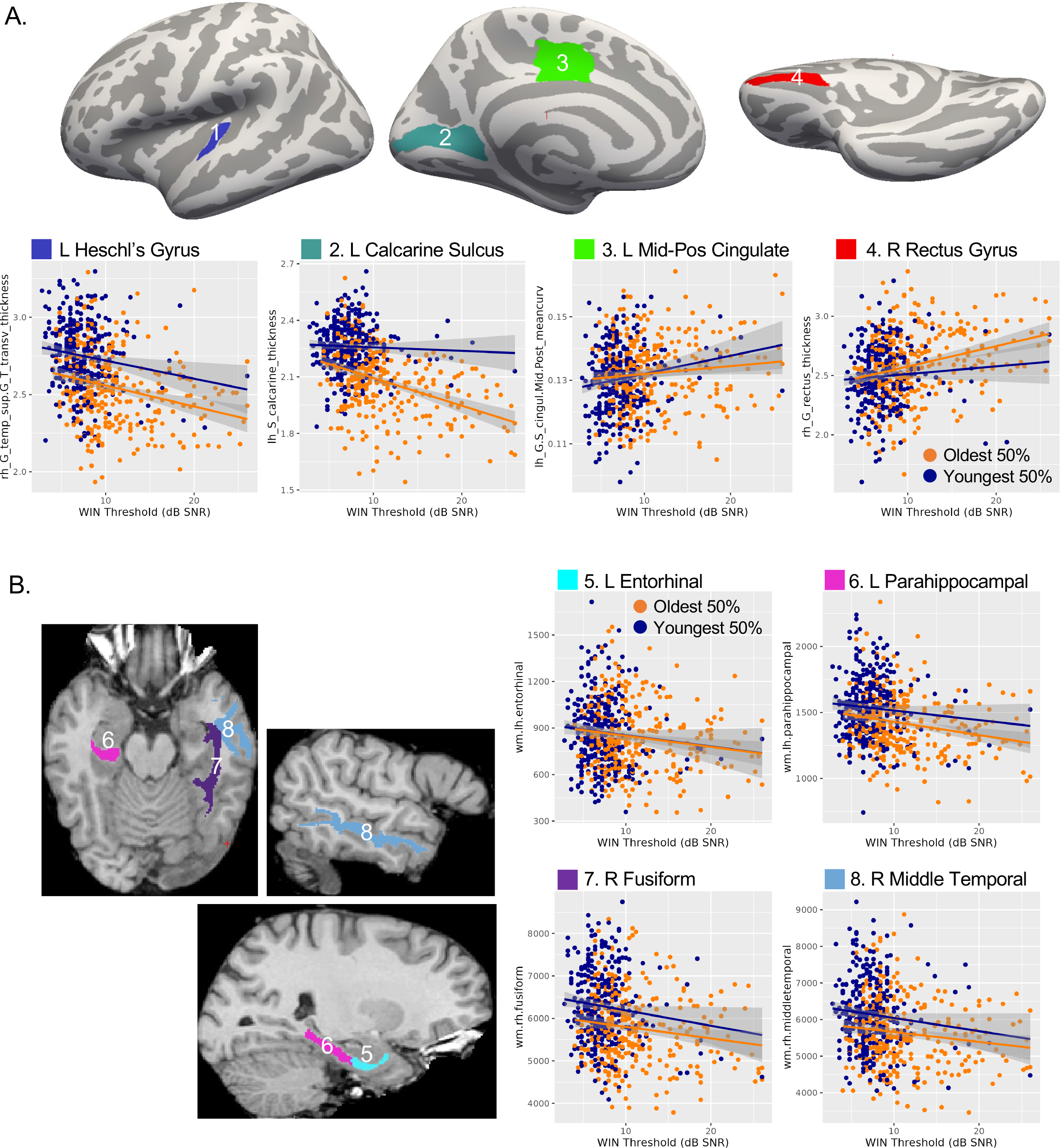
WIN threshold correlates with brain structure independently of age-related change. **A.** Regions with thickness or curvature significantly correlated with WIN threshold are displayed on a template cortical surface (top). Scatter plots display cortical thickness (mm) or mean curvature and WIN threshold for each volunteer, with linear regression lines shown for the top 50% oldest and 50% youngest ages. As in Figure 2, age was binarized for display only. **B.** Regions where volume significantly correlated with WIN threshold are displayed on a single subject at left. Scatter plots are displayed at right as in A. Abbreviations: L, left; R, right; Inf, inferior; Mid-Pos, Mid-posterior.

**Table 3.** Main Effects of Words in Noise Threshold on Brain Structure, p-fdr < 0.05.

| Region, Measure | effect | $\beta$ | $\beta$ SE | t | df | p | p<sub>FDR</sub> | partial r<sup>2</sup> |
| --- | --- | --- | --- | --- | --- | --- | --- | --- |
| L Heschl's gyrus, Thickness | WIN | -0.009 | 0.002 | -3.644 | 619 | 0.0003 | 0.037 | 0.021 |
|  | Age | -0.001 | 0.00006 | -11.162 | 619 | 0.00000 | 0.00000 | 0.168 |
| L Entorhinal White Matter, Volume | WIN | -8.530 | 2.414 | -3.533 | 619 | 0.0004 | 0.037 | 0.020 |
|  | Age | -0.022 | 0.059 | -0.371 | 619 | 0.711 | 0.746 | 0.000 |
| L Parahippocampal White Matter, Volume | WIN | -8.197 | 2.341 | -3.502 | 618 | 0.000 | 0.037 | 0.019 |
|  | Age | -0.340 | 0.058 | -5.904 | 618 | 0.00000 | 0.00000 | 0.053 |
| R Fusiform White Matter, Volume | WIN | -30.667 | 8.676 | -3.535 | 619 | 0.0004 | 0.037 | 0.020 |
|  | Age | -1.448 | 0.214 | -6.775 | 619 | 0.00000 | 0.00000 | 0.069 |
| R Middle Temporal White Matter, Volume | WIN | -32.812 | 9.433 | -3.478 | 619 | 0.001 | 0.037 | 0.019 |
|  | Age | -1.389 | 0.232 | -5.978 | 619 | 0.00000 | 0.00000 | 0.055 |
| L Mid Pos Cingulate Gyrus & Sulcus, Mean Curvature | WIN | 0.0004 | 0.0001 | 3.571 | 619 | 0.0004 | 0.037 | 0.020 |
|  | Age | 0.00000 | 0.00000 | -0.585 | 619 | 0.559 | 0.614 | 0.001 |
| L Calcarine Sulcus, Thickness | WIN | -0.006 | 0.002 | -3.713 | 619 | 0.0002 | 0.037 | 0.022 |
|  | Age | -0.001 | 0.00004 | -14.500 | 619 | 0.00000 | 0.00000 | 0.254 |
| R Rectus Gyrus, Thickness | WIN | 0.011 | 0.003 | 3.723 | 618 | 0.0002 | 0.037 | 0.022 |
|  | Age | 0.0003 | 0.0001 | 3.495 | 618 | 0.001 | 0.001 | 0.019 |

Note: Values listed as 0.00000 are less than 0.00001.

For completeness, we also report metrics exhibiting main effects of age controlling for hearing loss. Such effects were present across many measures, and can be reviewed in **Supplemental Table 2**.

### Exploratory analysis of auditory cortex

In exploratory analyses of auditory cortex, two Heschl’s gyrus metrics showed modest WIN-by-age interactions, including right hemisphere white matter and left hemisphere curvature. In both cases, age-related decreases were modestly less pronounced in volunteers with poorer hearing (puncorr < 0.05; **Table 4** top, **Figure 4A**). Several metrics exhibited modest main effect of WIN threshold (puncorr < 0.05, **Table 4** bottom, **Figure 4B**), including negative correlations with cortical thickness in right Heschl’s gyrus, the entire left temporal plane (i.e., Heschl’s gyrus, Heschl’s sulcus, planum temporale, planum polare) and left lateral superior temporal gyrus. White matter in right superior temporal cortex and bilateral temporal pole were also negatively correlated with WIN threshold independent of age, while left planum temporale curvature showed the opposite pattern (more curvature with poorer hearing).

**Figure 4.**
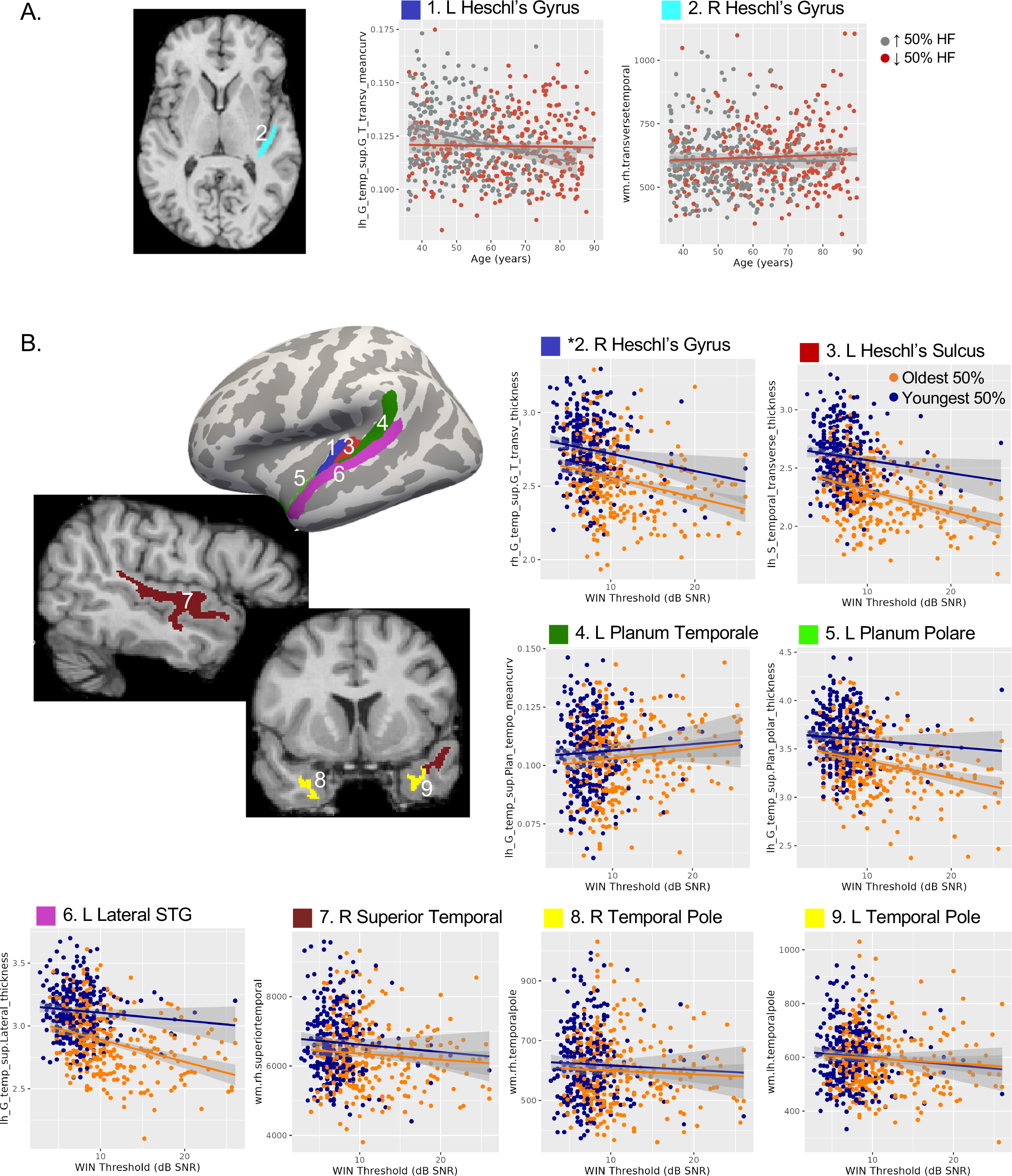
Exploratory analyses targeted auditory regions. **A.** A region showing WIN-by-age interaction puncorr < 0.05 are displayed a right, with scatterplots displayed as in Figure 2. **B.** Regions exhibiting main effects of WIN threshold independent of age puncorr < 0.05 are displayed at left on template cortical surface (thickness, mean curvature) and representative subject (volume). Scatter plots at right and bottom are displayed as in Figure 3. Abbreviations: L, left, R, right; STG, superior temporal gyrus.

**Table 4.** Auditory regions showing WIN-by-Age or main effects of WIN, p<0.05.

| Analysis | Region, Measure | effect | $\beta$ | $\beta$ SE | t | df | p | $p_{FDR}$ | partial $r^2$ |
| --- | --- | --- | --- | --- | --- | --- | --- | --- | --- |
| WIN*Age | R Heschl's Gyrus White Matter, Volume | WIN*Age | 0.017 | 0.007 | 2.518 | 617 | 0.012 | 0.308 | 0.010 |
|  |  | WIN | -15.281 | 5.780 | -2.644 | 617 | 0.008 | 0.686 | 0.011 |
|  |  | Age | -0.096 | 0.069 | -1.396 | 617 | 0.163 | 0.145 | 0.003 |
|  | L Heschl's Gyrus, Mean Curvature | WIN*Age | 0.00000 | 0.00000 | 2.835 | 618 | 0.005 | 0.185 | 0.013 |
|  |  | WIN | -0.002 | 0.001 | -2.468 | 618 | 0.014 | 0.495 | 0.010 |
|  |  | Age | 0.000 | 0.00001 | -4.442 | 618 | 0.00001 | 0.0002 | 0.031 |
| WIN | L Heschl's Sulcus, Thickness | WIN | -0.005 | 0.003 | -1.981 | 619 | 0.048 | 0.222 | 0.006 |
|  |  | Age | -0.001 | 0.000 | -15.747 | 619 | 0.000 | 0.000 | 0.286 |
|  | R Heschl's Gyrus, Thickness | WIN | -0.006 | 0.002 | -2.588 | 619 | 0.010 | 0.111 | 0.011 |
|  |  | Age | -0.001 | 0.000 | -10.475 | 619 | 0.00000 | 0.00000 | 0.151 |
|  | L Lateral STG, Thickness | WIN | -0.005 | 0.002 | -2.260 | 619 | 0.024 | 0.163 | 0.008 |
|  |  | Age | -0.001 | 0.000 | -14.635 | 619 | 0.000 | 0.000 | 0.257 |
|  | L Anterior STG, Thickness | WIN | -0.008 | 0.003 | -2.591 | 619 | 0.010 | 0.111 | 0.011 |
|  |  | Age | -0.001 | 0.000 | -9.725 | 619 | 0.00000 | 0.00000 | 0.133 |
|  | L Posterior STG, Curvature | WIN | 0.0004 | 0.0002 | 2.282 | 619 | 0.023 | 0.161 | 0.008 |
|  |  | Age | -0.00001 | 0.00000 | -2.050 | 619 | 0.041 | 0.061 | 0.007 |
|  | R STG White Matter, Volume | WIN | -30.912 | 9.694 | -3.189 | 619 | 0.002 | 0.063 | 0.016 |
|  |  | Age | -0.767 | 0.239 | -3.213 | 619 | 0.001 | 0.003 | 0.016 |
|  | L Temporal Pole White Matter, Volume | WIN | -3.281 | 1.175 | -2.791 | 619 | 0.005 | 0.092 | 0.012 |
|  |  | Age | -0.017 | 0.029 | -0.578 | 619 | 0.564 | 0.618 | 0.001 |
|  | R Temporal Pole White Matter, Volume | WIN | -2.390 | 1.201 | -1.990 | 619 | 0.047 | 0.221 | 0.006 |
|  |  | Age | -0.069 | 0.030 | -2.333 | 619 | 0.020 | 0.032 | 0.009 |

Note: Values listed as 0.00000 are less than 0.00001.

### Exploratory analysis of hearing and cognitive function

In the current sample, mean Montreal Cognitive Assessment (MoCA) total score was 26.37 (SD=2.54) and 225 of 623 volunteers had a score consistent with mild cognitive impairment (i.e., < 26, [Nasreddine et al., 2005]). For freesurfer metrics meeting statistical criterion pFDR<0.05 for either model described above, exploratory analyses measured a triple interaction between age, hearing loss, and total MoCA score (puncorr < 0.05; **Supplementary Table 3**). Of these, only left parahippocampal white matter was significant (t(613) = 2.18, p = 0.03, partial r^2^=0.008; **Figure 5**). Here, the slope of age-related volume loss was steepest in volunteers with poorer hearing and cognitive scores (i.e., higher WIN threshold and lower MoCA score). These interactions were not present for any other metric, though bilateral inferior lateral ventricles were both puncorr < 0.10.

**Figure 5.**
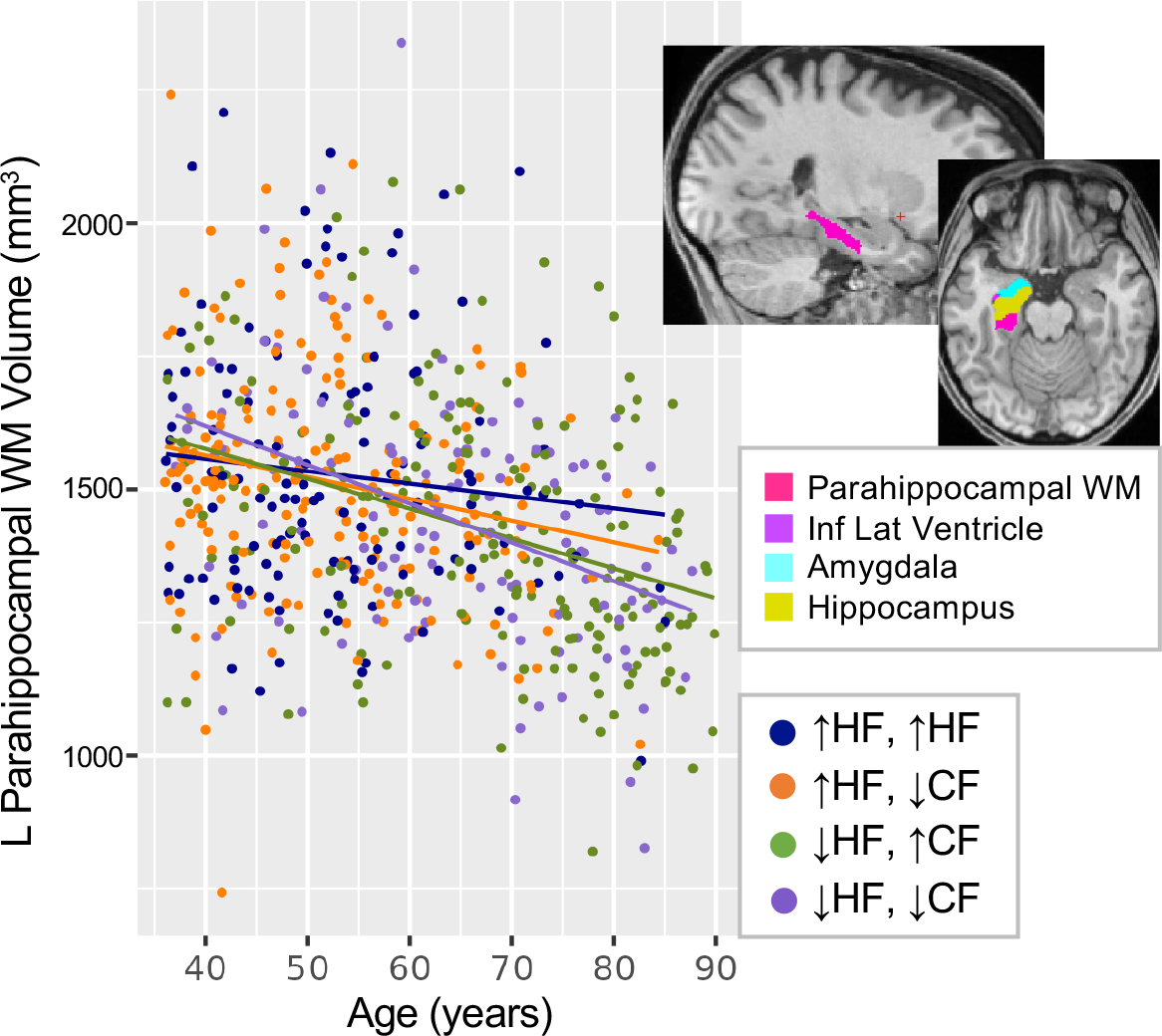
Exploratory analysis of cognitive function. Left parahippocampal white matter (pink in inset at upper right) showed a triple interaction between WIN threshold, age, and Montreal Cognitive Assessment (MoCA) score puncorr < 0.05. Scatter plot displays white matter volume (mm^3^) and age for each volunteer. For visualization only, WIN threshold and MoCA scores were binarized (top and bottom 50%), and regression lines are shown for each group. Abbreviations: HF, hearing function (WIN threshold); CF, cognitive function (MoCA score); ↑, top half of scores; ↓, bottom half of scores; L, left; WM, white matter; Inf, inferior; Lat, lateral.

## DISCUSSION

In the current study, we demonstrated that hearing function and age have both interacting and independent impacts on macro-anatomical brain features measured with MRI. In moderation analyses, poorer hearing function was associated with an accelerated pace of age-related differences, including increased ventricle size near medial temporal lobe structures. In primary auditory cortex and other sensory cortical regions, poorer hearing function correlated with reduced tissue content independent of age-related effects, suggesting that hearing loss may be linked to sensory cortical tissue loss at any age. Volunteers with poorer hearing and cognitive scores also tended to show steeper age-related reductions in left parahippocampal white matter. Taken together, these results suggest that hearing loss could be a modifiable risk factor in brain aging, particularly in medial temporal lobe structures affected in age-related neurodegenerative conditions like Alzheimer’s disease. However, longitudinal studies designed and powered to address the complexities of hearing loss are needed to directly assess causal relationships amongst hearing, brain aging, and age-related cognitive decline.

### Impact of hearing loss on brain aging

Many neuroimaging studies have reported relationships between hearing loss and brain structure and function. The current results replicate some of these findings, including correlations between Heschl’s gyrus and gyrus rectus gray matter and hearing scores while controlling for the effects of age [Eckert et al., 2012; Eckert et al., 2019; Husain et al., 2011; Koops et al., 2020; Lin et al., 2014; Melcher et al., 2013; Peelle et al., 2011]. We also report some novel findings, including an association between poorer hearing scores and reduced cortical thickness in primary visual cortex (i.e., calcarine sulcus), which could be interpreted as sensory deficits in one system impacting primary sensory cortex in other systems. This interpretation may also apply to findings in the gyrus rectus and mid-cingulate, which have been linked to olfaction [Rolls and Baylis, 1994] and somatosensation/pain [Sikes and Vogt, 1992; Wager et al., 2013], respectively. Although age-related differences were also apparent in these regions, hearing loss did not influence the rate of these trajectories. However, WIN- by-age interactions were present in left primary visual cortex and left visual association areas in lateral occipital cortex, suggesting that hearing loss may exacerbate age-related cortical thinning in these visual areas (or vice versa). WIN threshold also explained relatively more residual variance in lateral occipital cortex regions than age when controlling for the interaction term (**Table 2**). Taken together, these results suggest that hearing function may impact sensory cortices in all modalities, perhaps due to auditory deafferentation and/or loss of crossmodal cortico-cortical connections, and that these effects may be more pronounced in visual cortex in older adults. However, it is important to note that longitudinal studies are better suited to address causal relationships amongst these factors.

This point regarding causality is particularly salient when assessing whether our findings implicate hearing loss as a causal factor in anatomical changes in medial temporal lobe, which is heavily implicated in age-related cognitive impairment and dementias. In the current study, hearing function moderated age-related expansion of bilateral inferior lateral ventricles, located adjacent to entorhinal and parahippopcampal cortex, amygdala, and hippocampus. Ventricle expansion is a well-established biomarker of brain aging [Fujita et al., 2023; Irimia, 2021], and our current finding suggests that hearing loss associates with accelerated aging specifically in the medial temporal lobe. Given that the majority of volunteers in the current HCP Aging cohort likely had normal hearing (e.g., <6-10 dB SNR, [Humes, 2021; Leaver, 2024]) or mild loss, medial temporal lobe CSF could be particularly sensitive to the impacts of hearing function on age-related changes in this region. Indeed, pure-tone thresholds predicted overall ventricle expansion and white matter loss measured just ∼2.5 years later in volunteers aged ∼65 years old, suggesting that hearing loss may precede age-related ventricle expansion [Eckert et al., 2019].

We also noted correlations between WIN threshold and medial temporal lobe white matter in left entorhinal and parahippocampal cortex in the current study. This is consistent with previous studies reporting correlations between hearing loss and entorhinal and parahippocampal gray matter, as well as hippocampus and amygdala volume [Armstrong et al., 2019; Li et al., 2023; Rudner et al., 2019]. Notably, WIN threshold also correlated with gray matter volume in bilateral amygdala and hippocampus when controlling for age in our study, though these effects did not meet our strict correction for multiple comparisons (**Supplemental Table 2**). Though effects of age were also present in these structures, WIN-by-age interactions were not, suggesting that the pace of age-related changes in these structures were not impacted by hearing function in this cohort of healthy adults. Future studies including a wider range of hearing and cognitive function might be more sensitive to these relationships. Indeed, exploratory analyses indicated that age-related atrophy left parahippocampal white matter might be greatest in volunteers with both lower hearing and cognitive scores in our study. This is consistent with Li et al. 2023, who reported that gray matter in a small subregion of left parahippocampal gyrus mediated relationships between hearing and cognitive function in this same cohort [Li et al., 2023]. These effects are also compatible with Armstrong et al. 2019, who reported that hearing loss measured in at age ∼45 years predicted lower gray matter volume measured at age ∼65 in right hippocampus and left entorhinal cortex [Armstrong et al., 2019]. So, although the current study is cross-sectional, evidence from longitudinal studies suggest that hearing loss could be detectable before macro-anatomical tissue loss brain aging in medial temporal lobe regions implicated in age-related cognitive impairment.

However, though detectable hearing loss may precede brain aging measured with structural MRI, this does not necessarily mean that hearing loss causes brain aging. It is also possible that forms of brain aging not detectable on MRI (e.g., DNA methylation or other molecular changes [Horvath et al., 2012]) could cause and/or exacerbate the functional impacts of hearing loss. Auditory perception in a natural environment is exceedingly more complex than auditory sensation assessed with pure-tone audiometry, and relies on brain systems that analyze speech sounds, voice identity and inflection, separates sounds of interest from background noise, and so on [Peelle and Wingfield, 2016; Wayne and Johnsrude, 2015]. Therefore, it is possible that age-related atrophy and other changes in superior and/or medial temporal lobe make it more difficult to hear or compensate for hearing loss, thus worsening performance on pure-tone and/or words-in-noise detection thresholds during audiometric exams, or on cognitive exams. It is likely that causal relationships amongst hearing loss, brain aging, and age-related cognitive decline may not be unidirectional, and may instead mutually interact in different ways over time [Wayne and Johnsrude, 2015], and there are a number of other cogent reviews on this topic [Griffiths et al., 2020; Johnson et al., 2021; Wayne and Johnsrude, 2015; Whitson et al., 2018]. Yet, regardless of the precise causal mechanisms, the idea that early intervention with hearing aids or other assistive devices in midlife could delay brain aging remains compelling, especially if those interventions could delay the onset of age-related dementias or ameliorate their functional impact [Lin et al., 2023]. An alternate point of intervention could be to bolster attentional compensation strategies in hearing loss, which could be reflected by greater anterior cingulate cortex thickness in older volunteers with hearing loss in the current study [Pezzoli et al., 2024]. Longitudinal neuroimaging or other studies that measure brain aging in the same cohort over time are needed to assess both mechanistic causality, particularly those including comprehensive audiometric evaluations.

### WIN task performance as a potential measure of peripheral hearing loss

In the current study, we assumed that WIN task performance most likely reflected peripheral hearing loss in the HCP Aging cohort, where hearing loss was not exclusionary (to our understanding). Indeed, previous studies have noted correlations between WIN and pure-tone thresholds in typical populations [Fitzgerald et al., 2023; Holmes and Griffiths, 2019; Humes, 2021; Kam and Fu, 2020; Leaver, 2024; Vermiglio et al., 2020]. When administered without adjusting output volume to accommodate hearing loss in each volunteer, tablet-based WIN tasks can indeed be a quick, low-burden way of assessing hearing in large studies like the HCP, UK Biobank, and clinical trials [Leaver, 2024; Vermiglio et al., 2020], particularly when hearing is not the target of study.

Yet, it is widely understood that difficulty hearing in noisy environments can indicate central auditory dysfunction, independently of (or in conjunction with) peripheral hearing loss. Difficulty hearing in noise can occur in older adults without measurable peripheral hearing loss [Dubno et al., 2002; Helfer and Freyman, 2008; Schoof and Rosen, 2014], where tracking speech with competing talkers (vs. other types of noise) may be particularly affected [Helfer and Freyman, 2008; Rajan and Cainer, 2008; Schoof and Rosen, 2014; Tun et al., 2002]. In people with peripheral hearing loss, amplification does not always improve hearing in noise, though directionality settings may be under-utilized [Davidson et al., 2022]. However, it is unclear why such hearing in noise difficulties arise, and few therapies are available if amplification strategies fail. Therefore, there is a clear need to understand the brain bases of hearing in noise difficulties, and our study and others using commonly available datasets from the HCP, UK Biobank, ADNI, and others do not include comprehensive audiometry and are not able to address these nuances. So, although analyzing WIN task performance in these and similar datasets can improve our understanding of how hearing function impacts brain aging, large-scale studies combining full audiometry, neuroimaging, and other measures are needed.

One pattern of results noted in the current study may be relevant speech perception. When controlling for age, WIN threshold explained a significant amount of variance in structural metrics in the temporal lobe. Notably, effects in superior temporal regions tended to occur in gray matter in the left hemisphere, and in white matter in the right hemisphere. It is well-established that speech perception relies predominantly (though not exclusively) on left superior temporal regions [Leaver and Rauschecker, 2010; Scott and Johnsrude, 2003]. However, our results also suggest that right hemisphere white matter connections may also be important for typical speech perception and/or compensatory strategies in difficult hearing situations (e.g., using prosodic or timbre cues when decoding noisy speech). Given the limitations of the current study, it is difficult to say definitively that this pattern is the result of difficulty hearing speech in noise vs. peripheral hearing loss. However, it would be interesting to dissociate the impacts of central speech hearing difficulties and peripheral loss on brain structure and function in these populations in future studies [Holmes and Griffiths, 2019].

## Limitations

As with any study, there are limitations that should be considered when interpreting the current results. Perhaps most importantly, it is important to reiterate that the goal of the HCP Aging study was to characterize healthy brain aging, and so the current study was not designed *a priori* to study hearing loss or cognitive impairment. So, although the current data include a range of hearing and cognitive scores, a study that includes fuller variability on these measures with a more balanced number of volunteers with hearing loss and/or cognitive impairment might be more sensitive to the types of effects we sought to identify here. Similarly, though the Words in Noise task is very likely to approximate peripheral hearing loss in this sample, studies including full audiometric assessment are needed to dissociate contributions of central vs. peripheral hearing function. Despite these limitations, the current study and others like it provide evidence to support a role for hearing loss in brain aging in the medial temporal lobe elsewhere, motivating future studies of hearing loss to promote healthy brain aging.

## Conclusions

These findings provide evidence that age-related changes to brain morphometry are moderated by hearing function. In particular, poorer hearing correlates with age-related increased volume in bilateral inferior ventricles and with increased thinning in occipital structures. Additionally, even when controlling for age, WIN threshold explained variations in several temporal regions, including thinning of the left Heschl’s gyrus. These findings are consistent both with hearing-related changes to auditory structures, and with changes to brain structures associated with other sensory systems. Our results provide additional evidence linking age-related tissue loss in the left parahippocampal cortex with both hearing loss and poorer cognitive scores [Li et al., 2023], and going further by demonstrating that hearing loss may be a key driving factor in brain aging in this region. Taken together, these findings offer support for early interventions such as hearing aids to delay age-related changes to brain structures. Such interventions could prove especially valuable insofar as hearing loss has also been shown to be correlated with cognitive function [Lin et al., 2013; Stevenson et al., 2022; Whitson et al., 2018]. Future research in this area should focus on the causal relationship of these associations, particularly longitudinal studies that assess the progression of hearing loss and cognitive function in tandem with changes to brain morphometry. These future studies may also benefit from the use of functional neuroimaging modalities, such as arterial spin labelling MRI, which may be more sensitive to changes in persons at risk for cognitive impairment [Okonkwo et al., 2014]. Ideally, these studies should also include participants with clinically defined hearing loss or cognitive decline, in order to confirm these findings in target populations. Taken together, our results indicate the potential utility of such longitudinal studies in developing an understanding of the associations between hearing loss, brain structure, and cognitive decline, and that protecting hearing may be important for brain health.

## Acknowledgements

This work was supported by the National Institutes of Health under award number DC015880 to Dr. Leaver. This content is solely the responsibility of the authors and does not necessarily reflect the official views of the National Institutes of Health.

