## Supplemental Data for "Hearing function moderates age-related changes in brain morphometry in the HCP Aging cohort"

**Supplementary Table 1. WIN-by-Age Interaction Statistics For Metrics Showing Main Effects of WIN Threshold p-fdr < 0.05**

| Region, Measure | effect | beta | SE | t | df | p | fdr | paritalr2 |
|---|---|---|---|---|---|---|---|---|
| lh_G_temp_sup.G_T_transv_thickness | WIN*Age | 0.000 | 0.000 | -0.165 | 618 | 0.869 | 0.983 | 0.000 |
| lh_G_temp_sup.G_T_transv_thickness | WIN | -0.007 | 0.010 | -0.699 | 618 | 0.485 | 0.037 | 0.001 |
| lh_G_temp_sup.G_T_transv_thickness | Age | -0.001 | 0.000 | -5.320 | 618 | 0.000 | 0.000 | 0.044 |
| wm.lh.entorhinal | WIN*Age | -0.006 | 0.012 | -0.520 | 618 | 0.603 | 0.931 | 0.000 |
| wm.lh.entorhinal | WIN | -3.353 | 10.238 | -0.328 | 618 | 0.743 | 0.037 | 0.000 |
| wm.lh.entorhinal | Age | 0.033 | 0.121 | 0.272 | 618 | 0.786 | 0.746 | 0.000 |
| wm.lh.parahippocampal | WIN*Age | -0.002 | 0.012 | -0.190 | 617 | 0.850 | 0.983 | 0.000 |
| wm.lh.parahippocampal | WIN | -6.369 | 9.915 | -0.642 | 617 | 0.521 | 0.037 | 0.001 |
| wm.lh.parahippocampal | Age | -0.321 | 0.118 | -2.725 | 617 | 0.007 | 0.000 | 0.012 |
| wm.rh.fusiform | WIN*Age | 0.008 | 0.043 | 0.191 | 618 | 0.849 | 0.983 | 0.000 |
| wm.rh.fusiform | WIN | -37.498 | 36.797 | -1.019 | 618 | 0.309 | 0.037 | 0.002 |
| wm.rh.fusiform | Age | -1.521 | 0.437 | -3.483 | 618 | 0.001 | 0.000 | 0.019 |
| wm.rh.middletemporal | WIN*Age | 0.029 | 0.047 | 0.626 | 618 | 0.532 | 0.905 | 0.001 |
| wm.rh.middletemporal | WIN | -57.127 | 39.995 | -1.428 | 618 | 0.154 | 0.037 | 0.003 |
| wm.rh.middletemporal | Age | -1.648 | 0.475 | -3.472 | 618 | 0.001 | 0.000 | 0.019 |
| lh_S_calcarine_thickness | WIN*Age | 0.000 | 0.000 | -3.065 | 618 | 0.002 | 0.113 | 0.015 |
| lh_S_calcarine_thickness | WIN | 0.014 | 0.007 | 2.096 | 618 | 0.036 | 0.037 | 0.007 |
| lh_S_calcarine_thickness | Age | 0.000 | 0.000 | -4.479 | 618 | 0.000 | 0.000 | 0.031 |
| lh_G.S_cingul.Mid.Post_meancurv | WIN*Age | 0.000 | 0.000 | -1.528 | 618 | 0.127 | 0.679 | 0.004 |
| lh_G.S_cingul.Mid.Post_meancurv | WIN | 0.001 | 0.001 | 2.329 | 618 | 0.020 | 0.037 | 0.009 |
| lh_G.S_cingul.Mid.Post_meancurv | Age | 0.000 | 0.000 | 1.045 | 618 | 0.296 | 0.614 | 0.002 |
| rh_G_rectus_thickness | WIN*Age | 0.000 | 0.000 | 1.444 | 617 | 0.149 | 0.727 | 0.003 |
| rh_G_rectus_thickness | WIN | -0.007 | 0.013 | -0.523 | 617 | 0.601 | 0.037 | 0.000 |
| rh_G_rectus_thickness | Age | 0.000 | 0.000 | 0.458 | 617 | 0.647 | 0.001 | 0.000 |

**Supplementary Table 2. Structural Metrics Showing Main Effect of Age Threshold p-fdr < 0.05**

| Freesurfer Label | Metric | effect | beta | SE | t | df | p | fdr | paritalr2 |
|---|---|---|---|---|---|---|---|---|---|
| Left.Lateral.Ventricle | Volume | Age | 22.843 | 1.736 | 13.157 | 615 | 0.000 | 0.000 | 0.220 |
| Left.Inf.Lat.Vent | Volume | Age | 0.808 | 0.057 | 14.159 | 612 | 0.000 | 0.000 | 0.247 |
| Left.Cerebellum.White.Matter | Volume | Age | -3.802 | 0.438 | -8.689 | 618 | 0.000 | 0.000 | 0.109 |
| Left.Cerebellum.Cortex | Volume | Age | -7.304 | 1.352 | -5.402 | 618 | 0.000 | 0.000 | 0.045 |
| Left.Thalamus.Proper | Volume | Age | -2.040 | 0.171 | -11.943 | 617 | 0.000 | 0.000 | 0.188 |
| Left.Putamen | Volume | Age | -1.522 | 0.133 | -11.467 | 617 | 0.000 | 0.000 | 0.176 |
| Left.Pallidum | Volume | Age | -0.383 | 0.058 | -6.565 | 618 | 0.000 | 0.000 | 0.065 |
| Xthird.Ventricle | Volume | Age | 1.825 | 0.104 | 17.552 | 617 | 0.000 | 0.000 | 0.333 |
| Xfourth.Ventricle | Volume | Age | 0.555 | 0.132 | 4.198 | 616 | 0.000 | 0.000 | 0.028 |
| Left.Hippocampus | Volume | Age | -0.931 | 0.097 | -9.605 | 618 | 0.000 | 0.000 | 0.130 |
| Left.Amygdala | Volume | Age | -0.549 | 0.047 | -11.673 | 618 | 0.000 | 0.000 | 0.181 |
| CSF | Volume | Age | 0.527 | 0.067 | 7.879 | 614 | 0.000 | 0.000 | 0.092 |
| Left.Accumbens.area | Volume | Age | -0.354 | 0.022 | -16.219 | 618 | 0.000 | 0.000 | 0.299 |
| Left.VentralDC | Volume | Age | -0.856 | 0.106 | -8.054 | 618 | 0.000 | 0.000 | 0.095 |
| Left.choroid.plexus | Volume | Age | 0.510 | 0.047 | 10.902 | 618 | 0.000 | 0.000 | 0.161 |
| Right.Lateral.Ventricle | Volume | Age | 20.334 | 1.505 | 13.514 | 614 | 0.000 | 0.000 | 0.229 |
| Right.Inf.Lat.Vent | Volume | Age | 0.757 | 0.059 | 12.834 | 615 | 0.000 | 0.000 | 0.211 |
| Right.Cerebellum.White.Matter | Volume | Age | -3.382 | 0.436 | -7.765 | 617 | 0.000 | 0.000 | 0.089 |
| Right.Cerebellum.Cortex | Volume | Age | -7.212 | 1.394 | -5.175 | 618 | 0.000 | 0.000 | 0.042 |
| Right.Thalamus.Proper | Volume | Age | -1.805 | 0.165 | -10.928 | 618 | 0.000 | 0.000 | 0.162 |
| Right.Putamen | Volume | Age | -1.488 | 0.143 | -10.429 | 616 | 0.000 | 0.000 | 0.150 |
| Right.Pallidum | Volume | Age | -0.224 | 0.056 | -4.024 | 618 | 0.000 | 0.000 | 0.026 |
| Right.Hippocampus | Volume | Age | -0.946 | 0.101 | -9.364 | 618 | 0.000 | 0.000 | 0.124 |
| Right.Amygdala | Volume | Age | -0.499 | 0.053 | -9.386 | 618 | 0.000 | 0.000 | 0.125 |
| Right.Accumbens.area | Volume | Age | -0.239 | 0.023 | -10.632 | 618 | 0.000 | 0.000 | 0.155 |
| Right.VentralDC | Volume | Age | -0.959 | 0.103 | -9.277 | 618 | 0.000 | 0.000 | 0.122 |
| wm.lh.bankssts | Volume | Age | -0.602 | 0.154 | -3.907 | 619 | 0.000 | 0.000 | 0.024 |
| wm.lh.caudalanteriorcingulate | Volume | Age | -0.388 | 0.115 | -3.380 | 617 | 0.001 | 0.002 | 0.018 |
| wm.lh.caudalmiddlefrontal | Volume | Age | -0.689 | 0.253 | -2.717 | 618 | 0.007 | 0.012 | 0.012 |
| wm.lh.cuneus | Volume | Age | -0.405 | 0.116 | -3.493 | 619 | 0.001 | 0.001 | 0.019 |
| wm.lh.fusiform | Volume | Age | -1.586 | 0.227 | -6.982 | 618 | 0.000 | 0.000 | 0.073 |
| wm.lh.inferiorparietal | Volume | Age | -1.486 | 0.389 | -3.815 | 618 | 0.000 | 0.000 | 0.023 |
| wm.lh.inferiortemporal | Volume | Age | -1.197 | 0.265 | -4.511 | 619 | 0.000 | 0.000 | 0.032 |
| wm.lh.lateraloccipital | Volume | Age | -1.087 | 0.374 | -2.911 | 618 | 0.004 | 0.007 | 0.014 |
| wm.lh.lateralorbitofrontal | Volume | Age | -1.176 | 0.198 | -5.938 | 618 | 0.000 | 0.000 | 0.054 |
| wm.lh.lingual | Volume | Age | -0.936 | 0.205 | -4.572 | 618 | 0.000 | 0.000 | 0.033 |
| wm.lh.medialorbitofrontal | Volume | Age | -0.388 | 0.145 | -2.665 | 618 | 0.008 | 0.013 | 0.011 |
| wm.lh.middletemporal | Volume | Age | -1.331 | 0.224 | -5.951 | 619 | 0.000 | 0.000 | 0.054 |
| wm.lh.parahippocampal | Volume | Age | -0.340 | 0.058 | -5.904 | 618 | 0.000 | 0.000 | 0.053 |
| wm.lh.paracentral | Volume | Age | -0.486 | 0.142 | -3.415 | 618 | 0.001 | 0.001 | 0.019 |
| wm.lh.parsopercularis | Volume | Age | -0.675 | 0.158 | -4.263 | 618 | 0.000 | 0.000 | 0.029 |
| wm.lh.parsorbitalis | Volume | Age | -0.254 | 0.043 | -5.860 | 619 | 0.000 | 0.000 | 0.053 |
| wm.lh.parstriangularis | Volume | Age | -0.731 | 0.125 | -5.838 | 618 | 0.000 | 0.000 | 0.052 |
| wm.lh.pericalcarine | Volume | Age | -0.591 | 0.190 | -3.107 | 619 | 0.002 | 0.004 | 0.015 |
| wm.lh.posteriorcingulate | Volume | Age | -0.480 | 0.137 | -3.509 | 618 | 0.000 | 0.001 | 0.020 |
| wm.lh.precentral | Volume | Age | -1.214 | 0.400 | -3.033 | 618 | 0.003 | 0.005 | 0.015 |
| wm.lh.precuneus | Volume | Age | -0.892 | 0.347 | -2.573 | 618 | 0.010 | 0.017 | 0.011 |
| wm.lh.rostralmiddlefrontal | Volume | Age | -2.990 | 0.480 | -6.233 | 618 | 0.000 | 0.000 | 0.059 |
| wm.lh.superiorfrontal | Volume | Age | -3.500 | 0.606 | -5.780 | 618 | 0.000 | 0.000 | 0.051 |
| wm.lh.superiorparietal | Volume | Age | -1.660 | 0.425 | -3.906 | 618 | 0.000 | 0.000 | 0.024 |
| wm.lh.superiortemporal | Volume | Age | -1.248 | 0.288 | -4.338 | 618 | 0.000 | 0.000 | 0.030 |
| wm.lh.supramarginal | Volume | Age | -1.315 | 0.370 | -3.557 | 617 | 0.000 | 0.001 | 0.020 |

| | | | | | | | | | |
|---|---|---|---|---|---|---|---|---|---|
| wm.lh.frontalpole | Volume | Age | -0.050 | 0.012 | -4.149 | 618 | 0.000 | 0.000 | 0.027 |
| wm.rh.bankssts | Volume | Age | -0.483 | 0.138 | -3.492 | 619 | 0.001 | 0.001 | 0.019 |
| wm.rh.caudalanteriorcingulate | Volume | Age | -0.296 | 0.123 | -2.416 | 617 | 0.016 | 0.026 | 0.009 |
| wm.rh.cuneus | Volume | Age | -0.350 | 0.113 | -3.104 | 619 | 0.002 | 0.004 | 0.015 |
| wm.rh.fusiform | Volume | Age | -1.448 | 0.214 | -6.775 | 619 | 0.000 | 0.000 | 0.069 |
| wm.rh.inferiorparietal | Volume | Age | -1.336 | 0.436 | -3.065 | 619 | 0.002 | 0.004 | 0.015 |
| wm.rh.inferiortemporal | Volume | Age | -1.234 | 0.247 | -4.990 | 619 | 0.000 | 0.000 | 0.039 |
| wm.rh.lateraloccipital | Volume | Age | -1.469 | 0.384 | -3.825 | 618 | 0.000 | 0.000 | 0.023 |
| wm.rh.lateralorbitofrontal | Volume | Age | -1.063 | 0.219 | -4.847 | 619 | 0.000 | 0.000 | 0.037 |
| wm.rh.lingual | Volume | Age | -0.895 | 0.251 | -3.564 | 618 | 0.000 | 0.001 | 0.020 |
| wm.rh.medialorbitofrontal | Volume | Age | -0.524 | 0.120 | -4.351 | 619 | 0.000 | 0.000 | 0.030 |
| wm.rh.middletemporal | Volume | Age | -1.389 | 0.232 | -5.978 | 619 | 0.000 | 0.000 | 0.055 |
| wm.rh.parahippocampal | Volume | Age | -0.443 | 0.058 | -7.628 | 619 | 0.000 | 0.000 | 0.086 |
| wm.rh.paracentral | Volume | Age | -0.853 | 0.188 | -4.538 | 619 | 0.000 | 0.000 | 0.032 |
| wm.rh.parsopercularis | Volume | Age | -0.551 | 0.133 | -4.134 | 619 | 0.000 | 0.000 | 0.027 |
| wm.rh.parsorbitalis | Volume | Age | -0.254 | 0.051 | -5.014 | 619 | 0.000 | 0.000 | 0.039 |
| wm.rh.parstriangularis | Volume | Age | -0.804 | 0.142 | -5.673 | 618 | 0.000 | 0.000 | 0.050 |
| wm.rh.pericalcarine | Volume | Age | -0.578 | 0.197 | -2.933 | 619 | 0.003 | 0.006 | 0.014 |
| wm.rh.posteriorcingulate | Volume | Age | -0.631 | 0.137 | -4.612 | 619 | 0.000 | 0.000 | 0.033 |
| wm.rh.precentral | Volume | Age | -1.282 | 0.425 | -3.014 | 619 | 0.003 | 0.005 | 0.014 |
| wm.rh.precuneus | Volume | Age | -1.002 | 0.379 | -2.643 | 618 | 0.008 | 0.014 | 0.011 |
| wm.rh.rostralmiddlefrontal | Volume | Age | -2.583 | 0.534 | -4.839 | 618 | 0.000 | 0.000 | 0.037 |
| wm.rh.superiorfrontal | Volume | Age | -3.686 | 0.659 | -5.592 | 618 | 0.000 | 0.000 | 0.048 |
| wm.rh.superiorparietal | Volume | Age | -1.784 | 0.396 | -4.506 | 619 | 0.000 | 0.000 | 0.032 |
| wm.rh.superiortemporal | Volume | Age | -0.767 | 0.239 | -3.213 | 619 | 0.001 | 0.003 | 0.016 |
| wm.rh.frontalpole | Volume | Age | -0.044 | 0.016 | -2.855 | 619 | 0.004 | 0.008 | 0.013 |
| wm.rh.temporalpole | Volume | Age | -0.069 | 0.030 | -2.333 | 619 | 0.020 | 0.032 | 0.009 |
| lh_G.S_frontomargin_thickness | Cx Thickness | Age | 0.000 | 0.000 | -10.264 | 619 | 0.000 | 0.000 | 0.145 |
| lh_G.S_occipital_inf_thickness | Cx Thickness | Age | 0.000 | 0.000 | -7.404 | 618 | 0.000 | 0.000 | 0.081 |
| lh_G.S_paracentral_thickness | Cx Thickness | Age | 0.000 | 0.000 | -9.458 | 619 | 0.000 | 0.000 | 0.126 |
| lh_G.S_subcentral_thickness | Cx Thickness | Age | -0.001 | 0.000 | -15.421 | 619 | 0.000 | 0.000 | 0.278 |
| lh_G.S_transv_frontopol_thickness | Cx Thickness | Age | 0.000 | 0.000 | -8.847 | 619 | 0.000 | 0.000 | 0.112 |
| lh_G.S_cingul.Ant_thickness | Cx Thickness | Age | -0.001 | 0.000 | -11.993 | 619 | 0.000 | 0.000 | 0.189 |
| lh_G.S_cingul.Mid.Ant_thickness | Cx Thickness | Age | -0.001 | 0.000 | -17.021 | 619 | 0.000 | 0.000 | 0.319 |
| lh_G.S_cingul.Mid.Post_thickness | Cx Thickness | Age | -0.001 | 0.000 | -18.516 | 619 | 0.000 | 0.000 | 0.356 |
| lh_G_cingul.Post.dorsal_thickness | Cx Thickness | Age | -0.001 | 0.000 | -16.246 | 617 | 0.000 | 0.000 | 0.300 |
| lh_G_cingul.Post.ventral_thickness | Cx Thickness | Age | 0.000 | 0.000 | -6.678 | 619 | 0.000 | 0.000 | 0.067 |
| lh_G_cuneus_thickness | Cx Thickness | Age | 0.000 | 0.000 | -9.420 | 619 | 0.000 | 0.000 | 0.125 |
| lh_G_front_inf.Opercular_thickness | Cx Thickness | Age | -0.001 | 0.000 | -16.514 | 619 | 0.000 | 0.000 | 0.306 |
| lh_G_front_inf.Orbital_thickness | Cx Thickness | Age | -0.001 | 0.000 | -10.500 | 619 | 0.000 | 0.000 | 0.151 |
| lh_G_front_inf.Triangul_thickness | Cx Thickness | Age | -0.001 | 0.000 | -15.300 | 619 | 0.000 | 0.000 | 0.274 |
| lh_G_front_middle_thickness | Cx Thickness | Age | -0.001 | 0.000 | -17.612 | 619 | 0.000 | 0.000 | 0.334 |
| lh_G_front_sup_thickness | Cx Thickness | Age | -0.001 | 0.000 | -17.633 | 619 | 0.000 | 0.000 | 0.334 |
| lh_G_Ins_lg.S_cent_ins_thickness | Cx Thickness | Age | 0.000 | 0.000 | -6.652 | 619 | 0.000 | 0.000 | 0.067 |
| lh_G_insular_short_thickness | Cx Thickness | Age | -0.001 | 0.000 | -7.611 | 618 | 0.000 | 0.000 | 0.086 |
| lh_G_occipital_middle_thickness | Cx Thickness | Age | -0.001 | 0.000 | -12.642 | 619 | 0.000 | 0.000 | 0.205 |
| lh_G_occipital_sup_thickness | Cx Thickness | Age | 0.000 | 0.000 | -10.347 | 619 | 0.000 | 0.000 | 0.147 |
| lh_G_oc.temp_lat.fusifor_thickness | Cx Thickness | Age | -0.001 | 0.000 | -11.560 | 619 | 0.000 | 0.000 | 0.178 |
| lh_G_oc.temp_med.Lingual_thickness | Cx Thickness | Age | 0.000 | 0.000 | -7.676 | 619 | 0.000 | 0.000 | 0.087 |
| lh_G_oc.temp_med.Parahip_thickness | Cx Thickness | Age | 0.000 | 0.000 | -6.343 | 617 | 0.000 | 0.000 | 0.061 |
| lh_G_orbital_thickness | Cx Thickness | Age | 0.000 | 0.000 | -8.099 | 619 | 0.000 | 0.000 | 0.096 |
| lh_G_pariet_inf.Angular_thickness | Cx Thickness | Age | -0.001 | 0.000 | -17.838 | 619 | 0.000 | 0.000 | 0.340 |
| lh_G_pariet_inf.Supramar_thickness | Cx Thickness | Age | -0.001 | 0.000 | -16.055 | 619 | 0.000 | 0.000 | 0.294 |
| lh_G_parietal_sup_thickness | Cx Thickness | Age | -0.001 | 0.000 | -12.335 | 619 | 0.000 | 0.000 | 0.197 |

| | | | | | | | | | |
|---|---|---|---|---|---|---|---|---|---|
| lh_G_postcentral_thickness | Cx Thickness | Age | 0.000 | 0.000 | -11.648 | 619 | 0.000 | 0.000 | 0.180 |
| lh_G_precentral_thickness | Cx Thickness | Age | -0.001 | 0.000 | -16.665 | 619 | 0.000 | 0.000 | 0.310 |
| lh_G_precuneus_thickness | Cx Thickness | Age | -0.001 | 0.000 | -11.512 | 619 | 0.000 | 0.000 | 0.176 |
| lh_G_rectus_thickness | Cx Thickness | Age | 0.000 | 0.000 | 3.272 | 619 | 0.001 | 0.002 | 0.017 |
| lh_G_subcallosal_thickness | Cx Thickness | Age | -0.001 | 0.000 | -8.424 | 619 | 0.000 | 0.000 | 0.103 |
| lh_G_temp_sup.G_T_transv_thickness | Cx Thickness | Age | -0.001 | 0.000 | -11.162 | 619 | 0.000 | 0.000 | 0.168 |
| lh_G_temp_sup.Lateral_thickness | Cx Thickness | Age | -0.001 | 0.000 | -14.635 | 619 | 0.000 | 0.000 | 0.257 |
| lh_G_temp_sup.Plan_polar_thickness | Cx Thickness | Age | -0.001 | 0.000 | -9.725 | 619 | 0.000 | 0.000 | 0.133 |
| lh_G_temp_sup.Plan_tempo_thickness | Cx Thickness | Age | -0.001 | 0.000 | -14.291 | 619 | 0.000 | 0.000 | 0.248 |
| lh_G_temporal_inf_thickness | Cx Thickness | Age | 0.000 | 0.000 | -10.280 | 619 | 0.000 | 0.000 | 0.146 |
| lh_G_temporal_middle_thickness | Cx Thickness | Age | -0.001 | 0.000 | -15.655 | 619 | 0.000 | 0.000 | 0.284 |
| lh_Lat_Fis.ant.Horizont_thickness | Cx Thickness | Age | 0.000 | 0.000 | -8.308 | 618 | 0.000 | 0.000 | 0.100 |
| lh_Lat_Fis.ant.Vertical_thickness | Cx Thickness | Age | -0.001 | 0.000 | -12.634 | 619 | 0.000 | 0.000 | 0.205 |
| lh_Lat_Fis.post_thickness | Cx Thickness | Age | -0.001 | 0.000 | -18.294 | 619 | 0.000 | 0.000 | 0.351 |
| lh_Pole_occipital_thickness | Cx Thickness | Age | 0.000 | 0.000 | -5.875 | 619 | 0.000 | 0.000 | 0.053 |
| lh_Pole_temporal_thickness | Cx Thickness | Age | -0.001 | 0.000 | -10.289 | 619 | 0.000 | 0.000 | 0.146 |
| lh_S_calcarine_thickness | Cx Thickness | Age | -0.001 | 0.000 | -14.500 | 619 | 0.000 | 0.000 | 0.254 |
| lh_S_central_thickness | Cx Thickness | Age | -0.001 | 0.000 | -15.996 | 619 | 0.000 | 0.000 | 0.292 |
| lh_S_cingul.Marginalis_thickness | Cx Thickness | Age | -0.001 | 0.000 | -12.080 | 619 | 0.000 | 0.000 | 0.191 |
| lh_S_circular_insula_ant_thickness | Cx Thickness | Age | -0.001 | 0.000 | -12.607 | 619 | 0.000 | 0.000 | 0.204 |
| lh_S_circular_insula_inf_thickness | Cx Thickness | Age | -0.001 | 0.000 | -17.672 | 619 | 0.000 | 0.000 | 0.335 |
| lh_S_circular_insula_sup_thickness | Cx Thickness | Age | -0.001 | 0.000 | -20.594 | 619 | 0.000 | 0.000 | 0.407 |
| lh_S_collat_transv_ant_thickness | Cx Thickness | Age | -0.001 | 0.000 | -8.936 | 619 | 0.000 | 0.000 | 0.114 |
| lh_S_collat_transv_post_thickness | Cx Thickness | Age | 0.000 | 0.000 | -6.967 | 619 | 0.000 | 0.000 | 0.073 |
| lh_S_front_inf_thickness | Cx Thickness | Age | -0.001 | 0.000 | -17.733 | 619 | 0.000 | 0.000 | 0.337 |
| lh_S_front_middle_thickness | Cx Thickness | Age | -0.001 | 0.000 | -13.866 | 619 | 0.000 | 0.000 | 0.237 |
| lh_S_front_sup_thickness | Cx Thickness | Age | -0.001 | 0.000 | -18.638 | 619 | 0.000 | 0.000 | 0.359 |
| lh_S_interm_prim.Jensen_thickness | Cx Thickness | Age | -0.001 | 0.000 | -8.665 | 619 | 0.000 | 0.000 | 0.108 |
| lh_S_intrapariet.P_trans_thickness | Cx Thickness | Age | -0.001 | 0.000 | -15.586 | 619 | 0.000 | 0.000 | 0.282 |
| lh_S_oc_middle.Lunatus_thickness | Cx Thickness | Age | 0.000 | 0.000 | -12.260 | 618 | 0.000 | 0.000 | 0.196 |
| lh_S_oc_sup.transversal_thickness | Cx Thickness | Age | -0.001 | 0.000 | -12.591 | 618 | 0.000 | 0.000 | 0.204 |
| lh_S_occipital_ant_thickness | Cx Thickness | Age | -0.001 | 0.000 | -12.700 | 619 | 0.000 | 0.000 | 0.207 |
| lh_S_oc.temp_lat_thickness | Cx Thickness | Age | -0.001 | 0.000 | -10.394 | 619 | 0.000 | 0.000 | 0.149 |
| lh_S_oc.temp_med.Lingual_thickness | Cx Thickness | Age | -0.001 | 0.000 | -16.548 | 619 | 0.000 | 0.000 | 0.307 |
| lh_S_orbital_lateral_thickness | Cx Thickness | Age | -0.001 | 0.000 | -10.201 | 619 | 0.000 | 0.000 | 0.144 |
| lh_S_orbital_med.olfact_thickness | Cx Thickness | Age | 0.000 | 0.000 | -4.436 | 617 | 0.000 | 0.000 | 0.031 |
| lh_S_orbital.H_Shaped_thickness | Cx Thickness | Age | 0.000 | 0.000 | -10.027 | 619 | 0.000 | 0.000 | 0.140 |
| lh_S_parieto_occipital_thickness | Cx Thickness | Age | -0.001 | 0.000 | -17.289 | 619 | 0.000 | 0.000 | 0.326 |
| lh_S_pericallosal_thickness | Cx Thickness | Age | -0.001 | 0.000 | -11.316 | 619 | 0.000 | 0.000 | 0.171 |
| lh_S_postcentral_thickness | Cx Thickness | Age | -0.001 | 0.000 | -16.599 | 619 | 0.000 | 0.000 | 0.308 |
| lh_S_precentral.inf.part_thickness | Cx Thickness | Age | -0.001 | 0.000 | -18.153 | 619 | 0.000 | 0.000 | 0.347 |
| lh_S_precentral.sup.part_thickness | Cx Thickness | Age | -0.001 | 0.000 | -15.939 | 619 | 0.000 | 0.000 | 0.291 |
| lh_S_suborbital_thickness | Cx Thickness | Age | 0.000 | 0.000 | -4.174 | 618 | 0.000 | 0.000 | 0.027 |
| lh_S_subparietal_thickness | Cx Thickness | Age | -0.001 | 0.000 | -14.664 | 619 | 0.000 | 0.000 | 0.258 |
| lh_S_temporal_inf_thickness | Cx Thickness | Age | -0.001 | 0.000 | -14.015 | 619 | 0.000 | 0.000 | 0.241 |
| lh_S_temporal_sup_thickness | Cx Thickness | Age | -0.001 | 0.000 | -21.297 | 619 | 0.000 | 0.000 | 0.423 |
| lh_S_temporal_transverse_thickness | Cx Thickness | Age | -0.001 | 0.000 | -15.747 | 619 | 0.000 | 0.000 | 0.286 |
| lh_MeanThickness_thickness | Cx Thickness | Age | -0.001 | 0.000 | -22.273 | 619 | 0.000 | 0.000 | 0.445 |
| rh_G.S_frontomargin_thickness | Cx Thickness | Age | 0.000 | 0.000 | -8.622 | 619 | 0.000 | 0.000 | 0.107 |
| rh_G.S_occipital_inf_thickness | Cx Thickness | Age | 0.000 | 0.000 | -7.796 | 619 | 0.000 | 0.000 | 0.089 |
| rh_G.S_paracentral_thickness | Cx Thickness | Age | 0.000 | 0.000 | -10.102 | 619 | 0.000 | 0.000 | 0.142 |
| rh_G.S_subcentral_thickness | Cx Thickness | Age | -0.001 | 0.000 | -14.736 | 619 | 0.000 | 0.000 | 0.260 |
| rh_G.S_transv_frontopol_thickness | Cx Thickness | Age | 0.000 | 0.000 | -10.194 | 619 | 0.000 | 0.000 | 0.144 |
| rh_G.S_cingul.Ant_thickness | Cx Thickness | Age | 0.000 | 0.000 | -9.093 | 619 | 0.000 | 0.000 | 0.118 |

| | | | | | | | | | |
|---|---|---|---|---|---|---|---|---|---|
| rh_G.S_cingul.Mid.Ant_thickness | Cx Thickness | Age | -0.001 | 0.000 | -16.137 | 619 | 0.000 | 0.000 | 0.296 |
| rh_G.S_cingul.Mid.Post_thickness | Cx Thickness | Age | -0.001 | 0.000 | -18.501 | 619 | 0.000 | 0.000 | 0.356 |
| rh_G_cingul.Post.dorsal_thickness | Cx Thickness | Age | -0.001 | 0.000 | -17.956 | 619 | 0.000 | 0.000 | 0.342 |
| rh_G_cingul.Post.ventral_thickness | Cx Thickness | Age | 0.000 | 0.000 | -6.959 | 619 | 0.000 | 0.000 | 0.073 |
| rh_G_cuneus_thickness | Cx Thickness | Age | 0.000 | 0.000 | -9.462 | 619 | 0.000 | 0.000 | 0.126 |
| rh_G_front_inf.Opercular_thickness | Cx Thickness | Age | -0.001 | 0.000 | -17.671 | 619 | 0.000 | 0.000 | 0.335 |
| rh_G_front_inf.Orbital_thickness | Cx Thickness | Age | -0.001 | 0.000 | -11.475 | 619 | 0.000 | 0.000 | 0.175 |
| rh_G_front_inf.Triangul_thickness | Cx Thickness | Age | -0.001 | 0.000 | -14.398 | 619 | 0.000 | 0.000 | 0.251 |
| rh_G_front_middle_thickness | Cx Thickness | Age | -0.001 | 0.000 | -18.066 | 619 | 0.000 | 0.000 | 0.345 |
| rh_G_front_sup_thickness | Cx Thickness | Age | -0.001 | 0.000 | -16.465 | 619 | 0.000 | 0.000 | 0.305 |
| rh_G_Ins_lg.S_cent_ins_thickness | Cx Thickness | Age | 0.000 | 0.000 | -6.234 | 619 | 0.000 | 0.000 | 0.059 |
| rh_G_insular_short_thickness | Cx Thickness | Age | -0.001 | 0.000 | -7.722 | 619 | 0.000 | 0.000 | 0.088 |
| rh_G_occipital_middle_thickness | Cx Thickness | Age | -0.001 | 0.000 | -13.253 | 619 | 0.000 | 0.000 | 0.221 |
| rh_G_occipital_sup_thickness | Cx Thickness | Age | 0.000 | 0.000 | -10.468 | 619 | 0.000 | 0.000 | 0.150 |
| rh_G_oc.temp_lat.fusifor_thickness | Cx Thickness | Age | -0.001 | 0.000 | -11.721 | 619 | 0.000 | 0.000 | 0.182 |
| rh_G_oc.temp_med.Lingual_thickness | Cx Thickness | Age | 0.000 | 0.000 | -8.494 | 619 | 0.000 | 0.000 | 0.104 |
| rh_G_oc.temp_med.Parahip_thickness | Cx Thickness | Age | 0.000 | 0.000 | -6.799 | 619 | 0.000 | 0.000 | 0.069 |
| rh_G_orbital_thickness | Cx Thickness | Age | 0.000 | 0.000 | -6.989 | 619 | 0.000 | 0.000 | 0.073 |
| rh_G_pariet_inf.Angular_thickness | Cx Thickness | Age | -0.001 | 0.000 | -19.367 | 619 | 0.000 | 0.000 | 0.377 |
| rh_G_pariet_inf.Supramar_thickness | Cx Thickness | Age | -0.001 | 0.000 | -17.748 | 619 | 0.000 | 0.000 | 0.337 |
| rh_G_parietal_sup_thickness | Cx Thickness | Age | -0.001 | 0.000 | -13.211 | 619 | 0.000 | 0.000 | 0.220 |
| rh_G_postcentral_thickness | Cx Thickness | Age | -0.001 | 0.000 | -11.782 | 619 | 0.000 | 0.000 | 0.183 |
| rh_G_precentral_thickness | Cx Thickness | Age | -0.001 | 0.000 | -14.594 | 618 | 0.000 | 0.000 | 0.256 |
| rh_G_precuneus_thickness | Cx Thickness | Age | -0.001 | 0.000 | -11.399 | 619 | 0.000 | 0.000 | 0.173 |
| rh_G_rectus_thickness | Cx Thickness | Age | 0.000 | 0.000 | 3.495 | 618 | 0.001 | 0.001 | 0.019 |
| rh_G_subcallosal_thickness | Cx Thickness | Age | -0.001 | 0.000 | -4.761 | 619 | 0.000 | 0.000 | 0.035 |
| rh_G_temp_sup.G_T_transv_thickness | Cx Thickness | Age | -0.001 | 0.000 | -10.475 | 619 | 0.000 | 0.000 | 0.151 |
| rh_G_temp_sup.Lateral_thickness | Cx Thickness | Age | -0.001 | 0.000 | -15.289 | 619 | 0.000 | 0.000 | 0.274 |
| rh_G_temp_sup.Plan_polar_thickness | Cx Thickness | Age | -0.001 | 0.000 | -12.000 | 619 | 0.000 | 0.000 | 0.189 |
| rh_G_temp_sup.Plan_tempo_thickness | Cx Thickness | Age | -0.001 | 0.000 | -13.886 | 619 | 0.000 | 0.000 | 0.238 |
| rh_G_temporal_inf_thickness | Cx Thickness | Age | -0.001 | 0.000 | -12.116 | 619 | 0.000 | 0.000 | 0.192 |
| rh_G_temporal_middle_thickness | Cx Thickness | Age | -0.001 | 0.000 | -15.127 | 619 | 0.000 | 0.000 | 0.270 |
| rh_Lat_Fis.ant.Horizont_thickness | Cx Thickness | Age | -0.001 | 0.000 | -11.692 | 619 | 0.000 | 0.000 | 0.181 |
| rh_Lat_Fis.ant.Vertical_thickness | Cx Thickness | Age | -0.001 | 0.000 | -10.961 | 618 | 0.000 | 0.000 | 0.163 |
| rh_Lat_Fis.post_thickness | Cx Thickness | Age | -0.001 | 0.000 | -19.254 | 619 | 0.000 | 0.000 | 0.375 |
| rh_Pole_occipital_thickness | Cx Thickness | Age | 0.000 | 0.000 | -4.776 | 619 | 0.000 | 0.000 | 0.036 |
| rh_Pole_temporal_thickness | Cx Thickness | Age | -0.001 | 0.000 | -9.955 | 619 | 0.000 | 0.000 | 0.138 |
| rh_S_calcarine_thickness | Cx Thickness | Age | -0.001 | 0.000 | -15.394 | 619 | 0.000 | 0.000 | 0.277 |
| rh_S_central_thickness | Cx Thickness | Age | -0.001 | 0.000 | -15.848 | 619 | 0.000 | 0.000 | 0.289 |
| rh_S_cingul.Marginalis_thickness | Cx Thickness | Age | -0.001 | 0.000 | -13.181 | 618 | 0.000 | 0.000 | 0.219 |
| rh_S_circular_insula_ant_thickness | Cx Thickness | Age | -0.001 | 0.000 | -13.400 | 619 | 0.000 | 0.000 | 0.225 |
| rh_S_circular_insula_inf_thickness | Cx Thickness | Age | -0.001 | 0.000 | -18.216 | 619 | 0.000 | 0.000 | 0.349 |
| rh_S_circular_insula_sup_thickness | Cx Thickness | Age | -0.001 | 0.000 | -19.010 | 618 | 0.000 | 0.000 | 0.369 |
| rh_S_collat_transv_ant_thickness | Cx Thickness | Age | -0.001 | 0.000 | -10.705 | 619 | 0.000 | 0.000 | 0.156 |
| rh_S_collat_transv_post_thickness | Cx Thickness | Age | -0.001 | 0.000 | -10.388 | 619 | 0.000 | 0.000 | 0.148 |
| rh_S_front_inf_thickness | Cx Thickness | Age | -0.001 | 0.000 | -18.143 | 619 | 0.000 | 0.000 | 0.347 |
| rh_S_front_middle_thickness | Cx Thickness | Age | -0.001 | 0.000 | -14.355 | 619 | 0.000 | 0.000 | 0.250 |
| rh_S_front_sup_thickness | Cx Thickness | Age | -0.001 | 0.000 | -17.995 | 619 | 0.000 | 0.000 | 0.343 |
| rh_S_interm_prim.Jensen_thickness | Cx Thickness | Age | -0.001 | 0.000 | -13.575 | 619 | 0.000 | 0.000 | 0.229 |
| rh_S_intrapariet.P_trans_thickness | Cx Thickness | Age | -0.001 | 0.000 | -15.359 | 619 | 0.000 | 0.000 | 0.276 |
| rh_S_oc_middle.Lunatus_thickness | Cx Thickness | Age | 0.000 | 0.000 | -11.873 | 619 | 0.000 | 0.000 | 0.185 |
| rh_S_oc_sup.transversal_thickness | Cx Thickness | Age | -0.001 | 0.000 | -12.901 | 619 | 0.000 | 0.000 | 0.212 |
| rh_S_occipital_ant_thickness | Cx Thickness | Age | -0.001 | 0.000 | -12.972 | 619 | 0.000 | 0.000 | 0.214 |
| rh_S_oc.temp_lat_thickness | Cx Thickness | Age | -0.001 | 0.000 | -9.792 | 618 | 0.000 | 0.000 | 0.134 |

| | | | | | | | | | |
|---|---|---|---|---|---|---|---|---|---|
| rh_S_oc.temp_med.Lingual_thickness | Cx Thickness | Age | -0.001 | 0.000 | -16.877 | 619 | 0.000 | 0.000 | 0.315 |
| rh_S_orbital_lateral_thickness | Cx Thickness | Age | -0.001 | 0.000 | -12.691 | 619 | 0.000 | 0.000 | 0.206 |
| rh_S_orbital.H_Shaped_thickness | Cx Thickness | Age | 0.000 | 0.000 | -9.184 | 619 | 0.000 | 0.000 | 0.120 |
| rh_S_parieto_occipital_thickness | Cx Thickness | Age | -0.001 | 0.000 | -18.430 | 619 | 0.000 | 0.000 | 0.354 |
| rh_S_pericallosal_thickness | Cx Thickness | Age | -0.001 | 0.000 | -11.452 | 619 | 0.000 | 0.000 | 0.175 |
| rh_S_postcentral_thickness | Cx Thickness | Age | -0.001 | 0.000 | -16.303 | 619 | 0.000 | 0.000 | 0.300 |
| rh_S_precentral.inf.part_thickness | Cx Thickness | Age | -0.001 | 0.000 | -17.180 | 619 | 0.000 | 0.000 | 0.323 |
| rh_S_precentral.sup.part_thickness | Cx Thickness | Age | -0.001 | 0.000 | -15.049 | 619 | 0.000 | 0.000 | 0.268 |
| rh_S_subparietal_thickness | Cx Thickness | Age | -0.001 | 0.000 | -15.418 | 619 | 0.000 | 0.000 | 0.277 |
| rh_S_temporal_inf_thickness | Cx Thickness | Age | -0.001 | 0.000 | -14.364 | 619 | 0.000 | 0.000 | 0.250 |
| rh_S_temporal_sup_thickness | Cx Thickness | Age | -0.001 | 0.000 | -19.905 | 619 | 0.000 | 0.000 | 0.390 |
| rh_S_temporal_transverse_thickness | Cx Thickness | Age | -0.001 | 0.000 | -16.628 | 619 | 0.000 | 0.000 | 0.309 |
| rh_MeanThickness_thickness | Cx Thickness | Age | -0.001 | 0.000 | -22.108 | 619 | 0.000 | 0.000 | 0.441 |
| lh_G.S_cingul.Ant_area | Cx Surface Area | Age | -0.138 | 0.055 | -2.485 | 619 | 0.013 | 0.022 | 0.010 |
| lh_G.S_cingul.Mid.Post_area | Cx Surface Area | Age | -0.072 | 0.031 | -2.313 | 618 | 0.021 | 0.033 | 0.009 |
| lh_G_cingul.Post.dorsal_area | Cx Surface Area | Age | -0.052 | 0.020 | -2.681 | 619 | 0.008 | 0.013 | 0.011 |
| lh_G_cuneus_area | Cx Surface Area | Age | -0.167 | 0.055 | -3.025 | 619 | 0.003 | 0.005 | 0.015 |
| lh_G_front_inf.Triangul_area | Cx Surface Area | Age | -0.108 | 0.038 | -2.816 | 619 | 0.005 | 0.009 | 0.013 |
| lh_G_front_middle_area | Cx Surface Area | Age | -0.257 | 0.118 | -2.173 | 619 | 0.030 | 0.046 | 0.008 |
| lh_G_Ins_lg.S_cent_ins_area | Cx Surface Area | Age | -0.043 | 0.019 | -2.286 | 618 | 0.023 | 0.035 | 0.008 |
| lh_G_occipital_sup_area | Cx Surface Area | Age | -0.191 | 0.039 | -4.834 | 619 | 0.000 | 0.000 | 0.036 |
| lh_G_orbital_area | Cx Surface Area | Age | -0.172 | 0.047 | -3.658 | 619 | 0.000 | 0.001 | 0.021 |
| lh_G_pariet_inf.Angular_area | Cx Surface Area | Age | -0.238 | 0.067 | -3.552 | 619 | 0.000 | 0.001 | 0.020 |
| lh_G_subcallosal_area | Cx Surface Area | Age | 0.206 | 0.040 | 5.157 | 619 | 0.000 | 0.000 | 0.041 |
| lh_G_temp_sup.Plan_tempo_area | Cx Surface Area | Age | -0.121 | 0.036 | -3.331 | 618 | 0.001 | 0.002 | 0.018 |
| lh_G_temporal_inf_area | Cx Surface Area | Age | -0.251 | 0.073 | -3.421 | 619 | 0.001 | 0.001 | 0.019 |
| lh_Pole_occipital_area | Cx Surface Area | Age | -0.096 | 0.044 | -2.197 | 619 | 0.028 | 0.043 | 0.008 |
| lh_S_central_area | Cx Surface Area | Age | 0.300 | 0.061 | 4.901 | 618 | 0.000 | 0.000 | 0.037 |
| lh_S_collat_transv_ant_area | Cx Surface Area | Age | -0.090 | 0.034 | -2.678 | 619 | 0.008 | 0.013 | 0.011 |
| lh_S_interm_prim.Jensen_area | Cx Surface Area | Age | -0.103 | 0.033 | -3.097 | 619 | 0.002 | 0.004 | 0.015 |
| lh_S_oc.temp_lat_area | Cx Surface Area | Age | -0.107 | 0.033 | -3.204 | 619 | 0.001 | 0.003 | 0.016 |
| lh_S_oc.temp_med.Lingual_area | Cx Surface Area | Age | -0.216 | 0.045 | -4.778 | 618 | 0.000 | 0.000 | 0.036 |
| lh_S_orbital_lateral_area | Cx Surface Area | Age | -0.058 | 0.018 | -3.259 | 619 | 0.001 | 0.002 | 0.017 |
| lh_S_orbital_med.olfact_area | Cx Surface Area | Age | -0.063 | 0.016 | -3.909 | 619 | 0.000 | 0.000 | 0.024 |
| lh_S_orbital.H_Shaped_area | Cx Surface Area | Age | -0.086 | 0.033 | -2.609 | 619 | 0.009 | 0.016 | 0.011 |
| lh_S_postcentral_area | Cx Surface Area | Age | 0.269 | 0.084 | 3.204 | 619 | 0.001 | 0.003 | 0.016 |
| lh_S_subparietal_area | Cx Surface Area | Age | -0.096 | 0.036 | -2.690 | 618 | 0.007 | 0.013 | 0.012 |
| lh_S_temporal_sup_area | Cx Surface Area | Age | -0.380 | 0.129 | -2.955 | 619 | 0.003 | 0.006 | 0.014 |
| lh_WhiteSurfArea_area | Cx Surface Area | Age | -4.084 | 1.811 | -2.255 | 619 | 0.024 | 0.038 | 0.008 |
| rh_G.S_occipital_inf_area | Cx Surface Area | Age | -0.119 | 0.038 | -3.117 | 619 | 0.002 | 0.004 | 0.015 |
| rh_G_cuneus_area | Cx Surface Area | Age | -0.125 | 0.056 | -2.236 | 619 | 0.026 | 0.039 | 0.008 |
| rh_G_front_inf.Triangul_area | Cx Surface Area | Age | -0.117 | 0.037 | -3.164 | 618 | 0.002 | 0.003 | 0.016 |
| rh_G_occipital_sup_area | Cx Surface Area | Age | -0.169 | 0.042 | -3.995 | 619 | 0.000 | 0.000 | 0.025 |
| rh_G_oc.temp_med.Parahip_area | Cx Surface Area | Age | -0.105 | 0.043 | -2.455 | 619 | 0.014 | 0.023 | 0.010 |
| rh_G_orbital_area | Cx Surface Area | Age | -0.129 | 0.050 | -2.585 | 619 | 0.010 | 0.017 | 0.011 |
| rh_G_precentral_area | Cx Surface Area | Age | -0.133 | 0.055 | -2.412 | 619 | 0.016 | 0.026 | 0.009 |
| rh_G_temp_sup.Plan_tempo_area | Cx Surface Area | Age | -0.061 | 0.025 | -2.484 | 619 | 0.013 | 0.022 | 0.010 |
| rh_Pole_occipital_area | Cx Surface Area | Age | -0.241 | 0.075 | -3.207 | 619 | 0.001 | 0.003 | 0.016 |
| rh_S_central_area | Cx Surface Area | Age | 0.342 | 0.060 | 5.695 | 619 | 0.000 | 0.000 | 0.050 |
| rh_S_circular_insula_inf_area | Cx Surface Area | Age | 0.054 | 0.023 | 2.397 | 619 | 0.017 | 0.027 | 0.009 |
| rh_S_collat_transv_ant_area | Cx Surface Area | Age | -0.108 | 0.041 | -2.616 | 619 | 0.009 | 0.015 | 0.011 |
| rh_S_collat_transv_post_area | Cx Surface Area | Age | -0.047 | 0.020 | -2.325 | 619 | 0.020 | 0.032 | 0.009 |
| rh_S_occipital_ant_area | Cx Surface Area | Age | -0.079 | 0.033 | -2.389 | 619 | 0.017 | 0.027 | 0.009 |
| rh_S_oc.temp_lat_area | Cx Surface Area | Age | -0.097 | 0.042 | -2.317 | 619 | 0.021 | 0.033 | 0.009 |

| | | | | | | | | | |
|---|---|---|---|---|---|---|---|---|---|
| rh_S_oc.temp_med.Lingual_area | Cx Surface Area | Age | -0.222 | 0.050 | -4.434 | 619 | 0.000 | 0.000 | 0.031 |
| rh_S_orbital.H_Shaped_area | Cx Surface Area | Age | -0.112 | 0.033 | -3.339 | 619 | 0.001 | 0.002 | 0.018 |
| rh_S_postcentral_area | Cx Surface Area | Age | 0.294 | 0.083 | 3.547 | 619 | 0.000 | 0.001 | 0.020 |
| rh_S_subparietal_area | Cx Surface Area | Age | -0.106 | 0.043 | -2.450 | 619 | 0.015 | 0.024 | 0.010 |
| rh_S_temporal_inf_area | Cx Surface Area | Age | -0.094 | 0.044 | -2.168 | 619 | 0.031 | 0.046 | 0.008 |
| lh_G.S_frontomargin_meancurv | Cx Curvature | Age | 0.000 | 0.000 | 2.177 | 618 | 0.030 | 0.045 | 0.008 |
| lh_G.S_occipital_inf_meancurv | Cx Curvature | Age | 0.000 | 0.000 | 2.520 | 619 | 0.012 | 0.020 | 0.010 |
| lh_G.S_paracentral_meancurv | Cx Curvature | Age | 0.000 | 0.000 | 4.919 | 619 | 0.000 | 0.000 | 0.038 |
| lh_G.S_subcentral_meancurv | Cx Curvature | Age | 0.000 | 0.000 | 2.393 | 619 | 0.017 | 0.027 | 0.009 |
| lh_G.S_cingul.Mid.Ant_meancurv | Cx Curvature | Age | 0.000 | 0.000 | 2.195 | 619 | 0.029 | 0.043 | 0.008 |
| lh_G_cingul.Post.dorsal_meancurv | Cx Curvature | Age | 0.000 | 0.000 | 2.351 | 618 | 0.019 | 0.030 | 0.009 |
| lh_G_cuneus_meancurv | Cx Curvature | Age | 0.000 | 0.000 | -6.775 | 618 | 0.000 | 0.000 | 0.069 |
| lh_G_front_inf.Opercular_meancurv | Cx Curvature | Age | 0.000 | 0.000 | 3.543 | 618 | 0.000 | 0.001 | 0.020 |
| lh_G_front_middle_meancurv | Cx Curvature | Age | 0.000 | 0.000 | 3.339 | 618 | 0.001 | 0.002 | 0.018 |
| lh_G_front_sup_meancurv | Cx Curvature | Age | 0.000 | 0.000 | 4.012 | 618 | 0.000 | 0.000 | 0.025 |
| lh_G_insular_short_meancurv | Cx Curvature | Age | 0.000 | 0.000 | 7.325 | 619 | 0.000 | 0.000 | 0.080 |
| lh_G_occipital_middle_meancurv | Cx Curvature | Age | 0.000 | 0.000 | -2.964 | 619 | 0.003 | 0.006 | 0.014 |
| lh_G_occipital_sup_meancurv | Cx Curvature | Age | 0.000 | 0.000 | -7.272 | 619 | 0.000 | 0.000 | 0.079 |
| lh_G_oc.temp_lat.fusifor_meancurv | Cx Curvature | Age | 0.000 | 0.000 | 6.145 | 619 | 0.000 | 0.000 | 0.057 |
| lh_G_oc.temp_med.Lingual_meancurv | Cx Curvature | Age | 0.000 | 0.000 | -3.305 | 619 | 0.001 | 0.002 | 0.017 |
| lh_G_oc.temp_med.Parahip_meancurv | Cx Curvature | Age | 0.000 | 0.000 | 6.443 | 619 | 0.000 | 0.000 | 0.063 |
| lh_G_orbital_meancurv | Cx Curvature | Age | 0.000 | 0.000 | 5.863 | 619 | 0.000 | 0.000 | 0.053 |
| lh_G_pariet_inf.Supramar_meancurv | Cx Curvature | Age | 0.000 | 0.000 | 2.248 | 619 | 0.025 | 0.038 | 0.008 |
| lh_G_parietal_sup_meancurv | Cx Curvature | Age | 0.000 | 0.000 | -3.588 | 619 | 0.000 | 0.001 | 0.020 |
| lh_G_postcentral_meancurv | Cx Curvature | Age | 0.000 | 0.000 | -3.041 | 619 | 0.002 | 0.004 | 0.015 |
| lh_G_precentral_meancurv | Cx Curvature | Age | 0.000 | 0.000 | 6.465 | 617 | 0.000 | 0.000 | 0.063 |
| lh_G_rectus_meancurv | Cx Curvature | Age | 0.000 | 0.000 | 7.529 | 619 | 0.000 | 0.000 | 0.084 |
| lh_G_subcallosal_meancurv | Cx Curvature | Age | 0.000 | 0.000 | 4.941 | 619 | 0.000 | 0.000 | 0.038 |
| lh_G_temp_sup.G_T_transv_meancurv | Cx Curvature | Age | 0.000 | 0.000 | -4.000 | 619 | 0.000 | 0.000 | 0.025 |
| lh_G_temp_sup.Plan_polar_meancurv | Cx Curvature | Age | 0.000 | 0.000 | 2.302 | 619 | 0.022 | 0.034 | 0.008 |
| lh_G_temporal_inf_meancurv | Cx Curvature | Age | 0.000 | 0.000 | 7.088 | 619 | 0.000 | 0.000 | 0.075 |
| lh_G_temporal_middle_meancurv | Cx Curvature | Age | 0.000 | 0.000 | 7.907 | 618 | 0.000 | 0.000 | 0.092 |
| lh_Lat_Fis.ant.Horizont_meancurv | Cx Curvature | Age | 0.000 | 0.000 | 2.403 | 619 | 0.017 | 0.027 | 0.009 |
| lh_Lat_Fis.ant.Vertical_meancurv | Cx Curvature | Age | 0.000 | 0.000 | 3.059 | 617 | 0.002 | 0.004 | 0.015 |
| lh_Pole_temporal_meancurv | Cx Curvature | Age | 0.000 | 0.000 | 8.557 | 617 | 0.000 | 0.000 | 0.106 |
| lh_S_central_meancurv | Cx Curvature | Age | 0.000 | 0.000 | 6.001 | 618 | 0.000 | 0.000 | 0.055 |
| lh_S_cingul.Marginalis_meancurv | Cx Curvature | Age | 0.000 | 0.000 | -3.192 | 619 | 0.001 | 0.003 | 0.016 |
| lh_S_circular_insula_ant_meancurv | Cx Curvature | Age | 0.000 | 0.000 | 5.967 | 617 | 0.000 | 0.000 | 0.055 |
| lh_S_circular_insula_inf_meancurv | Cx Curvature | Age | 0.000 | 0.000 | -4.514 | 619 | 0.000 | 0.000 | 0.032 |
| lh_S_circular_insula_sup_meancurv | Cx Curvature | Age | 0.000 | 0.000 | 8.727 | 611 | 0.000 | 0.000 | 0.111 |
| lh_S_collat_transv_ant_meancurv | Cx Curvature | Age | 0.000 | 0.000 | 3.323 | 618 | 0.001 | 0.002 | 0.018 |
| lh_S_front_inf_meancurv | Cx Curvature | Age | 0.000 | 0.000 | 3.286 | 618 | 0.001 | 0.002 | 0.017 |
| lh_S_front_sup_meancurv | Cx Curvature | Age | 0.000 | 0.000 | 2.698 | 618 | 0.007 | 0.012 | 0.012 |
| lh_S_interm_prim.Jensen_meancurv | Cx Curvature | Age | 0.000 | 0.000 | 2.299 | 618 | 0.022 | 0.034 | 0.008 |
| lh_S_oc_middle.Lunatus_meancurv | Cx Curvature | Age | 0.000 | 0.000 | -2.274 | 619 | 0.023 | 0.036 | 0.008 |
| lh_S_oc.temp_lat_meancurv | Cx Curvature | Age | 0.000 | 0.000 | 2.694 | 619 | 0.007 | 0.012 | 0.012 |
| lh_S_oc.temp_med.Lingual_meancurv | Cx Curvature | Age | 0.000 | 0.000 | 5.001 | 618 | 0.000 | 0.000 | 0.039 |
| lh_S_orbital_med.olfact_meancurv | Cx Curvature | Age | 0.000 | 0.000 | 4.089 | 618 | 0.000 | 0.000 | 0.026 |
| lh_S_orbital.H_Shaped_meancurv | Cx Curvature | Age | 0.000 | 0.000 | 4.333 | 618 | 0.000 | 0.000 | 0.029 |
| lh_S_precentral.inf.part_meancurv | Cx Curvature | Age | 0.000 | 0.000 | 4.128 | 619 | 0.000 | 0.000 | 0.027 |
| lh_S_temporal_inf_meancurv | Cx Curvature | Age | 0.000 | 0.000 | 3.174 | 618 | 0.002 | 0.003 | 0.016 |
| rh_G.S_paracentral_meancurv | Cx Curvature | Age | 0.000 | 0.000 | 4.461 | 618 | 0.000 | 0.000 | 0.031 |
| rh_G.S_cingul.Ant_meancurv | Cx Curvature | Age | 0.000 | 0.000 | 4.011 | 618 | 0.000 | 0.000 | 0.025 |
| rh_G.S_cingul.Mid.Ant_meancurv | Cx Curvature | Age | 0.000 | 0.000 | 3.384 | 618 | 0.001 | 0.001 | 0.018 |

| | | | | | | | | | |
|---|---|---|---|---|---|---|---|---|---|
| rh_G.S_cingul.Mid.Post_meancurv | Cx Curvature | Age | 0.000 | 0.000 | 3.384 | 619 | 0.001 | 0.001 | 0.018 |
| rh_G_cingul.Post.dorsal_meancurv | Cx Curvature | Age | 0.000 | 0.000 | 4.668 | 617 | 0.000 | 0.000 | 0.034 |
| rh_G_cuneus_meancurv | Cx Curvature | Age | 0.000 | 0.000 | -4.680 | 619 | 0.000 | 0.000 | 0.034 |
| rh_G_front_inf.Opercular_meancurv | Cx Curvature | Age | 0.000 | 0.000 | 5.427 | 619 | 0.000 | 0.000 | 0.045 |
| rh_G_front_inf.Triangul_meancurv | Cx Curvature | Age | 0.000 | 0.000 | 2.491 | 619 | 0.013 | 0.022 | 0.010 |
| rh_G_front_middle_meancurv | Cx Curvature | Age | 0.000 | 0.000 | 2.316 | 618 | 0.021 | 0.033 | 0.009 |
| rh_G_front_sup_meancurv | Cx Curvature | Age | 0.000 | 0.000 | 4.864 | 618 | 0.000 | 0.000 | 0.037 |
| rh_G_insular_short_meancurv | Cx Curvature | Age | 0.000 | 0.000 | 5.286 | 618 | 0.000 | 0.000 | 0.043 |
| rh_G_occipital_sup_meancurv | Cx Curvature | Age | 0.000 | 0.000 | -8.362 | 619 | 0.000 | 0.000 | 0.101 |
| rh_G_oc.temp_lat.fusifor_meancurv | Cx Curvature | Age | 0.000 | 0.000 | 4.539 | 619 | 0.000 | 0.000 | 0.032 |
| rh_G_oc.temp_med.Parahip_meancurv | Cx Curvature | Age | 0.000 | 0.000 | 5.594 | 619 | 0.000 | 0.000 | 0.048 |
| rh_G_orbital_meancurv | Cx Curvature | Age | 0.000 | 0.000 | 8.147 | 619 | 0.000 | 0.000 | 0.097 |
| rh_G_pariet_inf.Angular_meancurv | Cx Curvature | Age | 0.000 | 0.000 | -3.517 | 619 | 0.000 | 0.001 | 0.020 |
| rh_G_parietal_sup_meancurv | Cx Curvature | Age | 0.000 | 0.000 | -8.967 | 619 | 0.000 | 0.000 | 0.115 |
| rh_G_precentral_meancurv | Cx Curvature | Age | 0.000 | 0.000 | 6.491 | 618 | 0.000 | 0.000 | 0.064 |
| rh_G_rectus_meancurv | Cx Curvature | Age | 0.000 | 0.000 | 6.410 | 618 | 0.000 | 0.000 | 0.062 |
| rh_G_subcallosal_meancurv | Cx Curvature | Age | 0.000 | 0.000 | 3.430 | 618 | 0.001 | 0.001 | 0.019 |
| rh_G_temp_sup.G_T_transv_meancurv | Cx Curvature | Age | 0.000 | 0.000 | -2.585 | 619 | 0.010 | 0.017 | 0.011 |
| rh_G_temporal_inf_meancurv | Cx Curvature | Age | 0.000 | 0.000 | 9.254 | 618 | 0.000 | 0.000 | 0.122 |
| rh_G_temporal_middle_meancurv | Cx Curvature | Age | 0.000 | 0.000 | 4.840 | 619 | 0.000 | 0.000 | 0.036 |
| rh_Lat_Fis.ant.Horizont_meancurv | Cx Curvature | Age | 0.000 | 0.000 | 3.347 | 619 | 0.001 | 0.002 | 0.018 |
| rh_Pole_temporal_meancurv | Cx Curvature | Age | 0.000 | 0.000 | 6.616 | 619 | 0.000 | 0.000 | 0.066 |
| rh_S_central_meancurv | Cx Curvature | Age | 0.000 | 0.000 | 6.203 | 618 | 0.000 | 0.000 | 0.059 |
| rh_S_circular_insula_ant_meancurv | Cx Curvature | Age | 0.000 | 0.000 | 5.251 | 618 | 0.000 | 0.000 | 0.043 |
| rh_S_circular_insula_inf_meancurv | Cx Curvature | Age | 0.000 | 0.000 | -3.943 | 619 | 0.000 | 0.000 | 0.025 |
| rh_S_circular_insula_sup_meancurv | Cx Curvature | Age | 0.000 | 0.000 | 6.956 | 615 | 0.000 | 0.000 | 0.073 |
| rh_S_collat_transv_ant_meancurv | Cx Curvature | Age | 0.000 | 0.000 | 2.596 | 619 | 0.010 | 0.016 | 0.011 |
| rh_S_front_inf_meancurv | Cx Curvature | Age | 0.000 | 0.000 | 2.708 | 619 | 0.007 | 0.012 | 0.012 |
| rh_S_interm_prim.Jensen_meancurv | Cx Curvature | Age | 0.000 | 0.000 | 2.410 | 618 | 0.016 | 0.026 | 0.009 |
| rh_S_oc_middle.Lunatus_meancurv | Cx Curvature | Age | 0.000 | 0.000 | -2.274 | 619 | 0.023 | 0.036 | 0.008 |
| rh_S_occipital_ant_meancurv | Cx Curvature | Age | 0.000 | 0.000 | 2.479 | 619 | 0.013 | 0.022 | 0.010 |
| rh_S_oc.temp_lat_meancurv | Cx Curvature | Age | 0.000 | 0.000 | 2.820 | 619 | 0.005 | 0.009 | 0.013 |
| rh_S_oc.temp_med.Lingual_meancurv | Cx Curvature | Age | 0.000 | 0.000 | 3.678 | 618 | 0.000 | 0.001 | 0.021 |
| rh_S_orbital_lateral_meancurv | Cx Curvature | Age | 0.000 | 0.000 | 2.374 | 619 | 0.018 | 0.028 | 0.009 |
| rh_S_orbital_med.olfact_meancurv | Cx Curvature | Age | 0.000 | 0.000 | 9.012 | 616 | 0.000 | 0.000 | 0.116 |
| rh_S_orbital.H_Shaped_meancurv | Cx Curvature | Age | 0.000 | 0.000 | 3.739 | 617 | 0.000 | 0.000 | 0.022 |
| rh_S_pericallosal_meancurv | Cx Curvature | Age | 0.000 | 0.000 | 3.095 | 619 | 0.002 | 0.004 | 0.015 |
| rh_S_precentral.inf.part_meancurv | Cx Curvature | Age | 0.000 | 0.000 | 5.272 | 618 | 0.000 | 0.000 | 0.043 |
| rh_S_precentral.sup.part_meancurv | Cx Curvature | Age | 0.000 | 0.000 | 4.300 | 619 | 0.000 | 0.000 | 0.029 |
| rh_S_temporal_inf_meancurv | Cx Curvature | Age | 0.000 | 0.000 | 4.143 | 619 | 0.000 | 0.000 | 0.027 |
| Left.Lateral.Ventricle | Volume | WIN | 65.601 | 70.327 | 0.933 | 615 | 0.351 | 0.636 | 0.001 |
| Left.Inf.Lat.Vent | Volume | WIN | 4.493 | 2.353 | 1.910 | 612 | 0.057 | 0.244 | 0.006 |
| ***Left.Cerebellum.White.Matter*** | ***Volume*** | ***WIN*** | ***-46.832*** | ***17.743*** | ***-2.639*** | ***618*** | ***0.009*** | ***0.111*** | ***0.011*** |
| Left.Cerebellum.Cortex | Volume | WIN | -94.862 | 54.826 | -1.730 | 618 | 0.084 | 0.289 | 0.005 |
| ***Left.Thalamus.Proper*** | ***Volume*** | ***WIN*** | ***-19.481*** | ***6.919*** | ***-2.816*** | ***617*** | ***0.005*** | ***0.091*** | ***0.013*** |
| Left.Putamen | Volume | WIN | -4.452 | 5.401 | -0.824 | 617 | 0.410 | 0.687 | 0.001 |
| ***Left.Pallidum*** | ***Volume*** | ***WIN*** | ***-5.429*** | ***2.364*** | ***-2.296*** | ***618*** | ***0.022*** | ***0.161*** | ***0.008*** |
| Xthird.Ventricle | Volume | WIN | 7.967 | 4.208 | 1.893 | 617 | 0.059 | 0.245 | 0.006 |
| Xfourth.Ventricle | Volume | WIN | 7.035 | 5.357 | 1.313 | 616 | 0.190 | 0.454 | 0.003 |
| ***Left.Hippocampus*** | ***Volume*** | ***WIN*** | ***-12.122*** | ***3.931*** | ***-3.083*** | ***618*** | ***0.002*** | ***0.076*** | ***0.015*** |
| ***Left.Amygdala*** | ***Volume*** | ***WIN*** | ***-6.359*** | ***1.905*** | ***-3.337*** | ***618*** | ***0.001*** | ***0.053*** | ***0.018*** |
| CSF | Volume | WIN | 1.880 | 2.716 | 0.692 | 614 | 0.489 | 0.744 | 0.001 |
| ***Left.Accumbens.area*** | ***Volume*** | ***WIN*** | ***-1.931*** | ***0.885*** | ***-2.181*** | ***618*** | ***0.030*** | ***0.180*** | ***0.008*** |
| ***Left.VentralDC*** | ***Volume*** | ***WIN*** | ***-10.694*** | ***4.309*** | ***-2.482*** | ***618*** | ***0.013*** | ***0.123*** | ***0.010*** |

| | | | | | | | | | |
|---|---|---|---|---|---|---|---|---|---|
| Left.choroid.plexus | Volume | WIN | 1.983 | 1.898 | 1.045 | 618 | 0.297 | 0.580 | 0.002 |
| Right.Lateral.Ventricle | Volume | WIN | 24.684 | 61.482 | 0.401 | 614 | 0.688 | 0.872 | 0.000 |
| ***Right.Inf.Lat.Vent*** | ***Volume*** | ***WIN*** | ***5.956*** | ***2.406*** | ***2.475*** | ***615*** | ***0.014*** | ***0.124*** | ***0.010*** |
| ***Right.Cerebellum.White.Matter*** | ***Volume*** | ***WIN*** | ***-35.797*** | ***17.639*** | ***-2.029*** | ***617*** | ***0.043*** | ***0.210*** | ***0.007*** |
| Right.Cerebellum.Cortex | Volume | WIN | -100.00 | 56.516 | -1.769 | 618 | 0.077 | 0.275 | 0.005 |
| ***Right.Thalamus.Proper*** | ***Volume*** | ***WIN*** | ***-15.261*** | ***6.698*** | ***-2.279*** | ***618*** | ***0.023*** | ***0.161*** | ***0.008*** |
| Right.Putamen | Volume | WIN | -1.572 | 5.764 | -0.273 | 616 | 0.785 | 0.917 | 0.000 |
| ***Right.Pallidum*** | ***Volume*** | ***WIN*** | ***-6.951*** | ***2.256*** | ***-3.081*** | ***618*** | ***0.002*** | ***0.076*** | ***0.015*** |
| ***Right.Hippocampus*** | ***Volume*** | ***WIN*** | ***-12.432*** | ***4.096*** | ***-3.036*** | ***618*** | ***0.003*** | ***0.076*** | ***0.015*** |
| ***Right.Amygdala*** | ***Volume*** | ***WIN*** | ***-6.434*** | ***2.157*** | ***-2.983*** | ***618*** | ***0.003*** | ***0.080*** | ***0.014*** |
| ***Right.Accumbens.area*** | ***Volume*** | ***WIN*** | ***-1.877*** | ***0.913*** | ***-2.057*** | ***618*** | ***0.040*** | ***0.206*** | ***0.007*** |
| ***Right.VentralDC*** | ***Volume*** | ***WIN*** | ***-9.746*** | ***4.193*** | ***-2.324*** | ***618*** | ***0.020*** | ***0.157*** | ***0.009*** |
| wm.lh.bankssts | Volume | WIN | -11.607 | 6.259 | -1.855 | 619 | 0.064 | 0.250 | 0.006 |
| wm.lh.caudalanteriorcingulate | Volume | WIN | 7.210 | 4.664 | 1.546 | 617 | 0.123 | 0.357 | 0.004 |
| wm.lh.caudalmiddlefrontal | Volume | WIN | 0.778 | 10.300 | 0.076 | 618 | 0.940 | 0.970 | 0.000 |
| wm.lh.cuneus | Volume | WIN | -3.007 | 4.705 | -0.639 | 619 | 0.523 | 0.780 | 0.001 |
| ***wm.lh.fusiform*** | ***Volume*** | ***WIN*** | ***-25.845*** | ***9.233*** | ***-2.799*** | ***618*** | ***0.005*** | ***0.092*** | ***0.013*** |
| wm.lh.inferiorparietal | Volume | WIN | -23.388 | 15.828 | -1.478 | 618 | 0.140 | 0.377 | 0.004 |
| ***wm.lh.inferiortemporal*** | ***Volume*** | ***WIN*** | ***-24.433*** | ***10.773*** | ***-2.268*** | ***619*** | ***0.024*** | ***0.163*** | ***0.008*** |
| wm.lh.lateraloccipital | Volume | WIN | -14.677 | 15.181 | -0.967 | 618 | 0.334 | 0.612 | 0.002 |
| ***wm.lh.lateralorbitofrontal*** | ***Volume*** | ***WIN*** | ***-23.922*** | ***8.048*** | ***-2.973*** | ***618*** | ***0.003*** | ***0.080*** | ***0.014*** |
| wm.lh.lingual | Volume | WIN | -13.786 | 8.321 | -1.657 | 618 | 0.098 | 0.317 | 0.004 |
| wm.lh.medialorbitofrontal | Volume | WIN | 3.663 | 5.913 | 0.619 | 618 | 0.536 | 0.795 | 0.001 |
| ***wm.lh.middletemporal*** | ***Volume*** | ***WIN*** | ***-22.646*** | ***9.079*** | ***-2.494*** | ***619*** | ***0.013*** | ***0.123*** | ***0.010*** |
| ***wm.lh.parahippocampal*** | ***Volume*** | ***WIN*** | ***-8.197*** | ***2.341*** | ***-3.502*** | ***618*** | ***0.000*** | ***0.037*** | ***0.019*** |
| ***wm.lh.paracentral*** | ***Volume*** | ***WIN*** | ***-16.918*** | ***5.787*** | ***-2.924*** | ***618*** | ***0.004*** | ***0.087*** | ***0.014*** |
| wm.lh.parsopercularis | Volume | WIN | -4.611 | 6.436 | -0.717 | 618 | 0.474 | 0.733 | 0.001 |
| wm.lh.parsorbitalis | Volume | WIN | -1.889 | 1.757 | -1.075 | 619 | 0.283 | 0.563 | 0.002 |
| wm.lh.parstriangularis | Volume | WIN | -4.612 | 5.087 | -0.907 | 618 | 0.365 | 0.647 | 0.001 |
| wm.lh.pericalcarine | Volume | WIN | -5.837 | 7.723 | -0.756 | 619 | 0.450 | 0.725 | 0.001 |
| ***wm.lh.posteriorcingulate*** | ***Volume*** | ***WIN*** | ***-10.949*** | ***5.565*** | ***-1.968*** | ***618*** | ***0.050*** | ***0.222*** | ***0.006*** |
| wm.lh.precentral | Volume | WIN | -19.345 | 16.273 | -1.189 | 618 | 0.235 | 0.503 | 0.002 |
| ***wm.lh.precuneus*** | ***Volume*** | ***WIN*** | ***-46.716*** | ***14.095*** | ***-3.314*** | ***618*** | ***0.001*** | ***0.053*** | ***0.017*** |
| wm.lh.rostralmiddlefrontal | Volume | WIN | -11.355 | 19.497 | -0.582 | 618 | 0.561 | 0.815 | 0.001 |
| ***wm.lh.superiorfrontal*** | ***Volume*** | ***WIN*** | ***-59.192*** | ***24.612*** | ***-2.405*** | ***618*** | ***0.016*** | ***0.136*** | ***0.009*** |
| ***wm.lh.superiorparietal*** | ***Volume*** | ***WIN*** | ***-35.569*** | ***17.268*** | ***-2.060*** | ***618*** | ***0.040*** | ***0.206*** | ***0.007*** |
| wm.lh.superiortemporal | Volume | WIN | -21.144 | 11.688 | -1.809 | 618 | 0.071 | 0.265 | 0.005 |
| wm.lh.supramarginal | Volume | WIN | -26.503 | 15.011 | -1.766 | 617 | 0.078 | 0.275 | 0.005 |
| wm.lh.frontalpole | Volume | WIN | -0.283 | 0.490 | -0.577 | 618 | 0.564 | 0.815 | 0.001 |
| ***wm.rh.bankssts*** | ***Volume*** | ***WIN*** | ***-11.257*** | ***5.610*** | ***-2.007*** | ***619*** | ***0.045*** | ***0.215*** | ***0.006*** |
| wm.rh.caudalanteriorcingulate | Volume | WIN | -9.004 | 4.977 | -1.809 | 617 | 0.071 | 0.265 | 0.005 |
| wm.rh.cuneus | Volume | WIN | -1.884 | 4.580 | -0.411 | 619 | 0.681 | 0.872 | 0.000 |
| ***wm.rh.fusiform*** | ***Volume*** | ***WIN*** | ***-30.667*** | ***8.676*** | ***-3.535*** | ***619*** | ***0.000*** | ***0.037*** | ***0.020*** |
| wm.rh.inferiorparietal | Volume | WIN | -8.473 | 17.700 | -0.479 | 619 | 0.632 | 0.846 | 0.000 |
| ***wm.rh.inferiortemporal*** | ***Volume*** | ***WIN*** | ***-22.961*** | ***10.036*** | ***-2.288*** | ***619*** | ***0.022*** | ***0.161*** | ***0.008*** |
| wm.rh.lateraloccipital | Volume | WIN | -13.936 | 15.572 | -0.895 | 618 | 0.371 | 0.652 | 0.001 |
| ***wm.rh.lateralorbitofrontal*** | ***Volume*** | ***WIN*** | ***-21.237*** | ***8.902*** | ***-2.386*** | ***619*** | ***0.017*** | ***0.141*** | ***0.009*** |
| wm.rh.lingual | Volume | WIN | -8.268 | 10.196 | -0.811 | 618 | 0.418 | 0.693 | 0.001 |
| wm.rh.medialorbitofrontal | Volume | WIN | -8.235 | 4.884 | -1.686 | 619 | 0.092 | 0.307 | 0.005 |
| ***wm.rh.middletemporal*** | ***Volume*** | ***WIN*** | ***-32.812*** | ***9.433*** | ***-3.478*** | ***619*** | ***0.001*** | ***0.037*** | ***0.019*** |
| wm.rh.parahippocampal | Volume | WIN | -4.439 | 2.356 | -1.884 | 619 | 0.060 | 0.246 | 0.006 |
| wm.rh.paracentral | Volume | WIN | -6.593 | 7.629 | -0.864 | 619 | 0.388 | 0.668 | 0.001 |
| wm.rh.parsopercularis | Volume | WIN | 1.491 | 5.413 | 0.276 | 619 | 0.783 | 0.917 | 0.000 |
| ***wm.rh.parsorbitalis*** | ***Volume*** | ***WIN*** | ***-4.222*** | ***2.054*** | ***-2.056*** | ***619*** | ***0.040*** | ***0.206*** | ***0.007*** |

| | | | | | | | | | |
|---|---|---|---|---|---|---|---|---|---|
| wm.rh.parstriangularis | Volume | WIN | -2.827 | 5.753 | -0.491 | 618 | 0.623 | 0.843 | 0.000 |
| wm.rh.pericalcarine | Volume | WIN | -10.437 | 8.008 | -1.303 | 619 | 0.193 | 0.458 | 0.003 |
| wm.rh.posteriorcingulate | Volume | WIN | -6.757 | 5.553 | -1.217 | 619 | 0.224 | 0.493 | 0.002 |
| wm.rh.precentral | Volume | WIN | -4.680 | 17.270 | -0.271 | 619 | 0.786 | 0.917 | 0.000 |
| ***wm.rh.precuneus*** | ***Volume*** | ***WIN*** | ***-43.654*** | ***15.397*** | ***-2.835*** | ***618*** | ***0.005*** | ***0.091*** | ***0.013*** |
| wm.rh.rostralmiddlefrontal | Volume | WIN | -26.155 | 21.637 | -1.209 | 618 | 0.227 | 0.494 | 0.002 |
| wm.rh.superiorfrontal | Volume | WIN | -37.094 | 26.756 | -1.386 | 618 | 0.166 | 0.423 | 0.003 |
| wm.rh.superiorparietal | Volume | WIN | -27.154 | 16.070 | -1.690 | 619 | 0.092 | 0.307 | 0.005 |
| ***wm.rh.superiortemporal*** | ***Volume*** | ***WIN*** | ***-30.912*** | ***9.694*** | ***-3.189*** | ***619*** | ***0.002*** | ***0.063*** | ***0.016*** |
| wm.rh.frontalpole | Volume | WIN | -0.153 | 0.630 | -0.242 | 619 | 0.808 | 0.930 | 0.000 |
| ***wm.rh.temporalpole*** | ***Volume*** | ***WIN*** | ***-2.390*** | ***1.201*** | ***-1.990*** | ***619*** | ***0.047*** | ***0.221*** | ***0.006*** |
| lh_G.S_frontomargin_thickness | Cx Thickness | WIN | 0.001 | 0.002 | 0.468 | 619 | 0.640 | 0.847 | 0.000 |
| ***lh_G.S_occipital_inf_thickness*** | ***Cx Thickness*** | ***WIN*** | ***-0.005*** | ***0.002*** | ***-2.718*** | ***618*** | ***0.007*** | ***0.095*** | ***0.012*** |
| ***lh_G.S_paracentral_thickness*** | ***Cx Thickness*** | ***WIN*** | ***-0.005*** | ***0.002*** | ***-2.298*** | ***619*** | ***0.022*** | ***0.161*** | ***0.008*** |
| lh_G.S_subcentral_thickness | Cx Thickness | WIN | -0.001 | 0.002 | -0.734 | 619 | 0.463 | 0.731 | 0.001 |
| lh_G.S_transv_frontopol_thickness | Cx Thickness | WIN | -0.002 | 0.002 | -1.021 | 619 | 0.308 | 0.587 | 0.002 |
| ***lh_G.S_cingul.Ant_thickness*** | ***Cx Thickness*** | ***WIN*** | ***0.004*** | ***0.002*** | ***2.179*** | ***619*** | ***0.030*** | ***0.180*** | ***0.008*** |
| lh_G.S_cingul.Mid.Ant_thickness | Cx Thickness | WIN | 0.001 | 0.002 | 0.463 | 619 | 0.643 | 0.847 | 0.000 |
| lh_G.S_cingul.Mid.Post_thickness | Cx Thickness | WIN | 0.001 | 0.002 | 0.346 | 619 | 0.730 | 0.889 | 0.000 |
| lh_G_cingul.Post.dorsal_thickness | Cx Thickness | WIN | -0.002 | 0.002 | -1.090 | 617 | 0.276 | 0.561 | 0.002 |
| lh_G_cingul.Post.ventral_thickness | Cx Thickness | WIN | -0.005 | 0.003 | -1.514 | 619 | 0.131 | 0.367 | 0.004 |
| ***lh_G_cuneus_thickness*** | ***Cx Thickness*** | ***WIN*** | ***-0.003*** | ***0.001*** | ***-2.174*** | ***619*** | ***0.030*** | ***0.180*** | ***0.008*** |
| lh_G_front_inf.Opercular_thickness | Cx Thickness | WIN | -0.001 | 0.002 | -0.705 | 619 | 0.481 | 0.742 | 0.001 |
| lh_G_front_inf.Orbital_thickness | Cx Thickness | WIN | -0.004 | 0.002 | -1.509 | 619 | 0.132 | 0.367 | 0.004 |
| lh_G_front_inf.Triangul_thickness | Cx Thickness | WIN | 0.000 | 0.002 | -0.223 | 619 | 0.824 | 0.935 | 0.000 |
| lh_G_front_middle_thickness | Cx Thickness | WIN | -0.001 | 0.002 | -0.535 | 619 | 0.593 | 0.837 | 0.000 |
| lh_G_front_sup_thickness | Cx Thickness | WIN | -0.002 | 0.002 | -0.886 | 619 | 0.376 | 0.658 | 0.001 |
| lh_G_Ins_lg.S_cent_ins_thickness | Cx Thickness | WIN | 0.000 | 0.003 | 0.004 | 619 | 0.997 | 0.999 | 0.000 |
| lh_G_insular_short_thickness | Cx Thickness | WIN | 0.000 | 0.003 | 0.126 | 618 | 0.899 | 0.961 | 0.000 |
| ***lh_G_occipital_middle_thickness*** | ***Cx Thickness*** | ***WIN*** | ***-0.005*** | ***0.002*** | ***-2.816*** | ***619*** | ***0.005*** | ***0.091*** | ***0.013*** |
| ***lh_G_occipital_sup_thickness*** | ***Cx Thickness*** | ***WIN*** | ***-0.005*** | ***0.002*** | ***-2.733*** | ***619*** | ***0.006*** | ***0.095*** | ***0.012*** |
| lh_G_oc.temp_lat.fusifor_thickness | Cx Thickness | WIN | -0.003 | 0.002 | -1.702 | 619 | 0.089 | 0.303 | 0.005 |
| ***lh_G_oc.temp_med.Lingual_thickness*** | ***Cx Thickness*** | ***WIN*** | ***-0.005*** | ***0.002*** | ***-3.280*** | ***619*** | ***0.001*** | ***0.055*** | ***0.017*** |
| ***lh_G_oc.temp_med.Parahip_thickness*** | ***Cx Thickness*** | ***WIN*** | ***-0.006*** | ***0.003*** | ***-2.162*** | ***617*** | ***0.031*** | ***0.180*** | ***0.008*** |
| lh_G_orbital_thickness | Cx Thickness | WIN | 0.001 | 0.002 | 0.694 | 619 | 0.488 | 0.744 | 0.001 |
| ***lh_G_pariet_inf.Angular_thickness*** | ***Cx Thickness*** | ***WIN*** | ***-0.005*** | ***0.002*** | ***-2.759*** | ***619*** | ***0.006*** | ***0.095*** | ***0.012*** |
| ***lh_G_pariet_inf.Supramar_thickness*** | ***Cx Thickness*** | ***WIN*** | ***-0.003*** | ***0.002*** | ***-2.068*** | ***619*** | ***0.039*** | ***0.206*** | ***0.007*** |
| lh_G_parietal_sup_thickness | Cx Thickness | WIN | -0.003 | 0.002 | -1.827 | 619 | 0.068 | 0.260 | 0.005 |
| lh_G_postcentral_thickness | Cx Thickness | WIN | 0.000 | 0.002 | -0.291 | 619 | 0.771 | 0.910 | 0.000 |
| ***lh_G_precentral_thickness*** | ***Cx Thickness*** | ***WIN*** | ***-0.005*** | ***0.002*** | ***-2.868*** | ***619*** | ***0.004*** | ***0.091*** | ***0.013*** |
| ***lh_G_precuneus_thickness*** | ***Cx Thickness*** | ***WIN*** | ***-0.005*** | ***0.002*** | ***-2.371*** | ***619*** | ***0.018*** | ***0.145*** | ***0.009*** |
| ***lh_G_rectus_thickness*** | ***Cx Thickness*** | ***WIN*** | ***0.006*** | ***0.002*** | ***2.404*** | ***619*** | ***0.016*** | ***0.136*** | ***0.009*** |
| lh_G_subcallosal_thickness | Cx Thickness | WIN | -0.003 | 0.004 | -0.692 | 619 | 0.489 | 0.744 | 0.001 |
| ***lh_G_temp_sup.G_T_transv_thickness*** | ***Cx Thickness*** | ***WIN*** | ***-0.009*** | ***0.002*** | ***-3.644*** | ***619*** | ***0.000*** | ***0.037*** | ***0.021*** |
| ***lh_G_temp_sup.Lateral_thickness*** | ***Cx Thickness*** | ***WIN*** | ***-0.005*** | ***0.002*** | ***-2.260*** | ***619*** | ***0.024*** | ***0.163*** | ***0.008*** |
| ***lh_G_temp_sup.Plan_polar_thickness*** | ***Cx Thickness*** | ***WIN*** | ***-0.008*** | ***0.003*** | ***-2.591*** | ***619*** | ***0.010*** | ***0.111*** | ***0.011*** |
| lh_G_temp_sup.Plan_tempo_thickness | Cx Thickness | WIN | 0.000 | 0.002 | 0.165 | 619 | 0.869 | 0.951 | 0.000 |
| lh_G_temporal_inf_thickness | Cx Thickness | WIN | -0.003 | 0.002 | -1.678 | 619 | 0.094 | 0.309 | 0.005 |
| ***lh_G_temporal_middle_thickness*** | ***Cx Thickness*** | ***WIN*** | ***-0.005*** | ***0.002*** | ***-2.748*** | ***619*** | ***0.006*** | ***0.095*** | ***0.012*** |
| lh_Lat_Fis.ant.Horizont_thickness | Cx Thickness | WIN | -0.002 | 0.002 | -0.979 | 618 | 0.328 | 0.608 | 0.002 |
| lh_Lat_Fis.ant.Vertical_thickness | Cx Thickness | WIN | -0.001 | 0.002 | -0.224 | 619 | 0.823 | 0.935 | 0.000 |
| lh_Lat_Fis.post_thickness | Cx Thickness | WIN | -0.002 | 0.002 | -1.135 | 619 | 0.257 | 0.534 | 0.002 |
| ***lh_Pole_occipital_thickness*** | ***Cx Thickness*** | ***WIN*** | ***-0.004*** | ***0.002*** | ***-2.235*** | ***619*** | ***0.026*** | ***0.169*** | ***0.008*** |
| lh_Pole_temporal_thickness | Cx Thickness | WIN | -0.003 | 0.002 | -1.546 | 619 | 0.123 | 0.357 | 0.004 |

| | | | | | | | | | |
|---|---|---|---|---|---|---|---|---|---|
| ***lh_S_calcarine_thickness*** | ***Cx Thickness*** | ***WIN*** | ***-0.006*** | ***0.002*** | ***-3.713*** | ***619*** | ***0.000*** | ***0.037*** | ***0.022*** |
| ***lh_S_central_thickness*** | ***Cx Thickness*** | ***WIN*** | ***-0.003*** | ***0.001*** | ***-2.112*** | ***619*** | ***0.035*** | ***0.195*** | ***0.007*** |
| lh_S_cingul.Marginalis_thickness | Cx Thickness | WIN | -0.003 | 0.002 | -1.516 | 619 | 0.130 | 0.367 | 0.004 |
| lh_S_circular_insula_ant_thickness | Cx Thickness | WIN | -0.001 | 0.002 | -0.410 | 619 | 0.682 | 0.872 | 0.000 |
| lh_S_circular_insula_inf_thickness | Cx Thickness | WIN | -0.002 | 0.002 | -1.219 | 619 | 0.223 | 0.493 | 0.002 |
| lh_S_circular_insula_sup_thickness | Cx Thickness | WIN | -0.001 | 0.002 | -0.420 | 619 | 0.675 | 0.871 | 0.000 |
| ***lh_S_collat_transv_ant_thickness*** | ***Cx Thickness*** | ***WIN*** | ***0.006*** | ***0.003*** | ***1.964*** | ***619*** | ***0.050*** | ***0.222*** | ***0.006*** |
| ***lh_S_collat_transv_post_thickness*** | ***Cx Thickness*** | ***WIN*** | ***-0.004*** | ***0.002*** | ***-2.196*** | ***619*** | ***0.028*** | ***0.179*** | ***0.008*** |
| lh_S_front_inf_thickness | Cx Thickness | WIN | 0.000 | 0.002 | 0.083 | 619 | 0.934 | 0.969 | 0.000 |
| lh_S_front_middle_thickness | Cx Thickness | WIN | 0.001 | 0.002 | 0.772 | 619 | 0.441 | 0.716 | 0.001 |
| lh_S_front_sup_thickness | Cx Thickness | WIN | 0.000 | 0.002 | -0.069 | 619 | 0.945 | 0.970 | 0.000 |
| lh_S_interm_prim.Jensen_thickness | Cx Thickness | WIN | -0.004 | 0.003 | -1.458 | 619 | 0.145 | 0.385 | 0.003 |
| ***lh_S_intrapariet.P_trans_thickness*** | ***Cx Thickness*** | ***WIN*** | ***-0.003*** | ***0.002*** | ***-2.141*** | ***619*** | ***0.033*** | ***0.186*** | ***0.007*** |
| ***lh_S_oc_middle.Lunatus_thickness*** | ***Cx Thickness*** | ***WIN*** | ***-0.004*** | ***0.001*** | ***-2.717*** | ***618*** | ***0.007*** | ***0.095*** | ***0.012*** |
| ***lh_S_oc_sup.transversal_thickness*** | ***Cx Thickness*** | ***WIN*** | ***-0.004*** | ***0.002*** | ***-2.561*** | ***618*** | ***0.011*** | ***0.111*** | ***0.010*** |
| ***lh_S_occipital_ant_thickness*** | ***Cx Thickness*** | ***WIN*** | ***-0.005*** | ***0.002*** | ***-2.562*** | ***619*** | ***0.011*** | ***0.111*** | ***0.010*** |
| ***lh_S_oc.temp_lat_thickness*** | ***Cx Thickness*** | ***WIN*** | ***-0.005*** | ***0.002*** | ***-2.242*** | ***619*** | ***0.025*** | ***0.169*** | ***0.008*** |
| lh_S_oc.temp_med.Lingual_thickness | Cx Thickness | WIN | -0.003 | 0.002 | -1.889 | 619 | 0.059 | 0.246 | 0.006 |
| lh_S_orbital_lateral_thickness | Cx Thickness | WIN | -0.002 | 0.002 | -0.763 | 619 | 0.446 | 0.722 | 0.001 |
| lh_S_orbital_med.olfact_thickness | Cx Thickness | WIN | 0.004 | 0.002 | 1.855 | 617 | 0.064 | 0.250 | 0.006 |
| lh_S_orbital.H_Shaped_thickness | Cx Thickness | WIN | 0.003 | 0.002 | 1.617 | 619 | 0.106 | 0.330 | 0.004 |
| lh_S_parieto_occipital_thickness | Cx Thickness | WIN | -0.003 | 0.002 | -1.647 | 619 | 0.100 | 0.318 | 0.004 |
| lh_S_pericallosal_thickness | Cx Thickness | WIN | 0.000 | 0.003 | -0.188 | 619 | 0.851 | 0.947 | 0.000 |
| lh_S_postcentral_thickness | Cx Thickness | WIN | -0.002 | 0.002 | -1.181 | 619 | 0.238 | 0.508 | 0.002 |
| lh_S_precentral.inf.part_thickness | Cx Thickness | WIN | 0.000 | 0.002 | 0.226 | 619 | 0.821 | 0.935 | 0.000 |
| lh_S_precentral.sup.part_thickness | Cx Thickness | WIN | -0.003 | 0.002 | -1.804 | 619 | 0.072 | 0.266 | 0.005 |
| lh_S_suborbital_thickness | Cx Thickness | WIN | 0.002 | 0.002 | 0.719 | 618 | 0.472 | 0.733 | 0.001 |
| ***lh_S_subparietal_thickness*** | ***Cx Thickness*** | ***WIN*** | ***-0.005*** | ***0.002*** | ***-2.458*** | ***619*** | ***0.014*** | ***0.128*** | ***0.010*** |
| lh_S_temporal_inf_thickness | Cx Thickness | WIN | -0.001 | 0.002 | -0.845 | 619 | 0.399 | 0.678 | 0.001 |
| ***lh_S_temporal_sup_thickness*** | ***Cx Thickness*** | ***WIN*** | ***-0.003*** | ***0.001*** | ***-2.083*** | ***619*** | ***0.038*** | ***0.203*** | ***0.007*** |
| ***lh_S_temporal_transverse_thickness*** | ***Cx Thickness*** | ***WIN*** | ***-0.005*** | ***0.003*** | ***-1.981*** | ***619*** | ***0.048*** | ***0.222*** | ***0.006*** |
| ***lh_MeanThickness_thickness*** | ***Cx Thickness*** | ***WIN*** | ***-0.002*** | ***0.001*** | ***-2.205*** | ***619*** | ***0.028*** | ***0.176*** | ***0.008*** |
| rh_G.S_frontomargin_thickness | Cx Thickness | WIN | 0.002 | 0.002 | 0.837 | 619 | 0.403 | 0.683 | 0.001 |
| rh_G.S_occipital_inf_thickness | Cx Thickness | WIN | -0.003 | 0.002 | -1.591 | 619 | 0.112 | 0.343 | 0.004 |
| ***rh_G.S_paracentral_thickness*** | ***Cx Thickness*** | ***WIN*** | ***-0.004*** | ***0.002*** | ***-2.328*** | ***619*** | ***0.020*** | ***0.157*** | ***0.009*** |
| rh_G.S_subcentral_thickness | Cx Thickness | WIN | -0.001 | 0.002 | -0.530 | 619 | 0.596 | 0.838 | 0.000 |
| rh_G.S_transv_frontopol_thickness | Cx Thickness | WIN | 0.001 | 0.002 | 0.754 | 619 | 0.451 | 0.725 | 0.001 |
| rh_G.S_cingul.Ant_thickness | Cx Thickness | WIN | 0.004 | 0.002 | 1.860 | 619 | 0.063 | 0.250 | 0.006 |
| rh_G.S_cingul.Mid.Ant_thickness | Cx Thickness | WIN | 0.000 | 0.002 | 0.146 | 619 | 0.884 | 0.958 | 0.000 |
| rh_G.S_cingul.Mid.Post_thickness | Cx Thickness | WIN | 0.001 | 0.002 | 0.781 | 619 | 0.435 | 0.711 | 0.001 |
| rh_G_cingul.Post.dorsal_thickness | Cx Thickness | WIN | 0.002 | 0.002 | 1.067 | 619 | 0.286 | 0.564 | 0.002 |
| rh_G_cingul.Post.ventral_thickness | Cx Thickness | WIN | -0.002 | 0.003 | -0.805 | 619 | 0.421 | 0.695 | 0.001 |
| rh_G_cuneus_thickness | Cx Thickness | WIN | -0.002 | 0.001 | -1.373 | 619 | 0.170 | 0.429 | 0.003 |
| rh_G_front_inf.Opercular_thickness | Cx Thickness | WIN | 0.001 | 0.002 | 0.659 | 619 | 0.510 | 0.765 | 0.001 |
| rh_G_front_inf.Orbital_thickness | Cx Thickness | WIN | -0.001 | 0.003 | -0.384 | 619 | 0.701 | 0.882 | 0.000 |
| rh_G_front_inf.Triangul_thickness | Cx Thickness | WIN | 0.000 | 0.002 | 0.097 | 619 | 0.923 | 0.966 | 0.000 |
| rh_G_front_middle_thickness | Cx Thickness | WIN | -0.002 | 0.002 | -0.967 | 619 | 0.334 | 0.612 | 0.002 |
| rh_G_front_sup_thickness | Cx Thickness | WIN | 0.000 | 0.002 | -0.271 | 619 | 0.787 | 0.917 | 0.000 |
| rh_G_Ins_lg.S_cent_ins_thickness | Cx Thickness | WIN | 0.001 | 0.003 | 0.307 | 619 | 0.759 | 0.902 | 0.000 |
| rh_G_insular_short_thickness | Cx Thickness | WIN | -0.002 | 0.003 | -0.698 | 619 | 0.486 | 0.744 | 0.001 |
| ***rh_G_occipital_middle_thickness*** | ***Cx Thickness*** | ***WIN*** | ***-0.003*** | ***0.002*** | ***-2.006*** | ***619*** | ***0.045*** | ***0.215*** | ***0.006*** |
| ***rh_G_occipital_sup_thickness*** | ***Cx Thickness*** | ***WIN*** | ***-0.005*** | ***0.002*** | ***-2.486*** | ***619*** | ***0.013*** | ***0.123*** | ***0.010*** |
| rh_G_oc.temp_lat.fusifor_thickness | Cx Thickness | WIN | -0.003 | 0.002 | -1.649 | 619 | 0.100 | 0.318 | 0.004 |
| rh_G_oc.temp_med.Lingual_thickness | Cx Thickness | WIN | -0.002 | 0.002 | -1.350 | 619 | 0.178 | 0.443 | 0.003 |

| | | | | | | | | | |
|---|---|---|---|---|---|---|---|---|---|
| rh_G_oc.temp_med.Parahip_thickness | Cx Thickness | WIN | -0.003 | 0.003 | -1.025 | 619 | 0.306 | 0.587 | 0.002 |
| rh_G_orbital_thickness | Cx Thickness | WIN | 0.002 | 0.002 | 1.166 | 619 | 0.244 | 0.516 | 0.002 |
| rh_G_pariet_inf.Angular_thickness | Cx Thickness | WIN | -0.003 | 0.002 | -1.586 | 619 | 0.113 | 0.343 | 0.004 |
| rh_G_pariet_inf.Supramar_thickness | Cx Thickness | WIN | -0.001 | 0.002 | -0.517 | 619 | 0.605 | 0.843 | 0.000 |
| rh_G_parietal_sup_thickness | Cx Thickness | WIN | -0.003 | 0.002 | -1.448 | 619 | 0.148 | 0.387 | 0.003 |
| rh_G_postcentral_thickness | Cx Thickness | WIN | -0.001 | 0.002 | -0.595 | 619 | 0.552 | 0.810 | 0.001 |
| rh_G_precentral_thickness | Cx Thickness | WIN | -0.004 | 0.002 | -1.752 | 618 | 0.080 | 0.281 | 0.005 |
| ***rh_G_precuneus_thickness*** | ***Cx Thickness*** | ***WIN*** | ***-0.004*** | ***0.002*** | ***-2.027*** | ***619*** | ***0.043*** | ***0.210*** | ***0.007*** |
| ***rh_G_rectus_thickness*** | ***Cx Thickness*** | ***WIN*** | ***0.011*** | ***0.003*** | ***3.723*** | ***618*** | ***0.000*** | ***0.037*** | ***0.022*** |
| rh_G_subcallosal_thickness | Cx Thickness | WIN | -0.003 | 0.006 | -0.496 | 619 | 0.620 | 0.843 | 0.000 |
| ***rh_G_temp_sup.G_T_transv_thickness*** | ***Cx Thickness*** | ***WIN*** | ***-0.006*** | ***0.002*** | ***-2.588*** | ***619*** | ***0.010*** | ***0.111*** | ***0.011*** |
| rh_G_temp_sup.Lateral_thickness | Cx Thickness | WIN | -0.002 | 0.002 | -0.718 | 619 | 0.473 | 0.733 | 0.001 |
| rh_G_temp_sup.Plan_polar_thickness | Cx Thickness | WIN | -0.003 | 0.003 | -1.079 | 619 | 0.281 | 0.562 | 0.002 |
| rh_G_temp_sup.Plan_tempo_thickness | Cx Thickness | WIN | 0.001 | 0.002 | 0.241 | 619 | 0.809 | 0.930 | 0.000 |
| rh_G_temporal_inf_thickness | Cx Thickness | WIN | -0.001 | 0.002 | -0.604 | 619 | 0.546 | 0.806 | 0.001 |
| rh_G_temporal_middle_thickness | Cx Thickness | WIN | -0.003 | 0.002 | -1.351 | 619 | 0.177 | 0.443 | 0.003 |
| rh_Lat_Fis.ant.Horizont_thickness | Cx Thickness | WIN | 0.000 | 0.002 | -0.048 | 619 | 0.962 | 0.978 | 0.000 |
| rh_Lat_Fis.ant.Vertical_thickness | Cx Thickness | WIN | 0.000 | 0.002 | -0.014 | 618 | 0.989 | 0.998 | 0.000 |
| rh_Lat_Fis.post_thickness | Cx Thickness | WIN | -0.001 | 0.002 | -0.341 | 619 | 0.733 | 0.889 | 0.000 |
| ***rh_Pole_occipital_thickness*** | ***Cx Thickness*** | ***WIN*** | ***-0.004*** | ***0.002*** | ***-2.842*** | ***619*** | ***0.005*** | ***0.091*** | ***0.013*** |
| ***rh_Pole_temporal_thickness*** | ***Cx Thickness*** | ***WIN*** | ***-0.004*** | ***0.002*** | ***-2.054*** | ***619*** | ***0.040*** | ***0.206*** | ***0.007*** |
| rh_S_calcarine_thickness | Cx Thickness | WIN | -0.002 | 0.002 | -1.468 | 619 | 0.143 | 0.380 | 0.003 |
| rh_S_central_thickness | Cx Thickness | WIN | -0.002 | 0.001 | -1.535 | 619 | 0.125 | 0.358 | 0.004 |
| rh_S_cingul.Marginalis_thickness | Cx Thickness | WIN | -0.001 | 0.002 | -0.695 | 618 | 0.488 | 0.744 | 0.001 |
| rh_S_circular_insula_ant_thickness | Cx Thickness | WIN | 0.004 | 0.002 | 1.792 | 619 | 0.074 | 0.270 | 0.005 |
| rh_S_circular_insula_inf_thickness | Cx Thickness | WIN | -0.001 | 0.002 | -0.464 | 619 | 0.643 | 0.847 | 0.000 |
| rh_S_circular_insula_sup_thickness | Cx Thickness | WIN | 0.002 | 0.002 | 0.978 | 618 | 0.328 | 0.608 | 0.002 |
| rh_S_collat_transv_ant_thickness | Cx Thickness | WIN | -0.001 | 0.003 | -0.234 | 619 | 0.815 | 0.933 | 0.000 |
| rh_S_collat_transv_post_thickness | Cx Thickness | WIN | -0.002 | 0.002 | -1.029 | 619 | 0.304 | 0.586 | 0.002 |
| rh_S_front_inf_thickness | Cx Thickness | WIN | 0.000 | 0.002 | 0.128 | 619 | 0.898 | 0.961 | 0.000 |
| rh_S_front_middle_thickness | Cx Thickness | WIN | 0.003 | 0.002 | 1.542 | 619 | 0.124 | 0.357 | 0.004 |
| rh_S_front_sup_thickness | Cx Thickness | WIN | 0.001 | 0.002 | 0.342 | 619 | 0.733 | 0.889 | 0.000 |
| rh_S_interm_prim.Jensen_thickness | Cx Thickness | WIN | 0.000 | 0.002 | -0.154 | 619 | 0.878 | 0.955 | 0.000 |
| rh_S_intrapariet.P_trans_thickness | Cx Thickness | WIN | -0.002 | 0.002 | -1.275 | 619 | 0.203 | 0.473 | 0.003 |
| ***rh_S_oc_middle.Lunatus_thickness*** | ***Cx Thickness*** | ***WIN*** | ***-0.005*** | ***0.002*** | ***-2.744*** | ***619*** | ***0.006*** | ***0.095*** | ***0.012*** |
| rh_S_oc_sup.transversal_thickness | Cx Thickness | WIN | -0.002 | 0.002 | -1.425 | 619 | 0.155 | 0.399 | 0.003 |
| rh_S_occipital_ant_thickness | Cx Thickness | WIN | -0.003 | 0.002 | -1.776 | 619 | 0.076 | 0.274 | 0.005 |
| ***rh_S_oc.temp_lat_thickness*** | ***Cx Thickness*** | ***WIN*** | ***-0.005*** | ***0.002*** | ***-2.220*** | ***618*** | ***0.027*** | ***0.172*** | ***0.008*** |
| rh_S_oc.temp_med.Lingual_thickness | Cx Thickness | WIN | 0.000 | 0.002 | -0.210 | 619 | 0.834 | 0.939 | 0.000 |
| rh_S_orbital_lateral_thickness | Cx Thickness | WIN | 0.000 | 0.002 | 0.117 | 619 | 0.907 | 0.961 | 0.000 |
| rh_S_orbital.H_Shaped_thickness | Cx Thickness | WIN | 0.003 | 0.002 | 1.546 | 619 | 0.123 | 0.357 | 0.004 |
| rh_S_parieto_occipital_thickness | Cx Thickness | WIN | 0.000 | 0.002 | -0.118 | 619 | 0.906 | 0.961 | 0.000 |
| rh_S_pericallosal_thickness | Cx Thickness | WIN | 0.001 | 0.003 | 0.565 | 619 | 0.573 | 0.823 | 0.001 |
| ***rh_S_postcentral_thickness*** | ***Cx Thickness*** | ***WIN*** | ***-0.003*** | ***0.002*** | ***-2.017*** | ***619*** | ***0.044*** | ***0.213*** | ***0.007*** |
| rh_S_precentral.inf.part_thickness | Cx Thickness | WIN | 0.002 | 0.002 | 1.507 | 619 | 0.132 | 0.367 | 0.004 |
| rh_S_precentral.sup.part_thickness | Cx Thickness | WIN | -0.001 | 0.002 | -0.690 | 619 | 0.491 | 0.744 | 0.001 |
| rh_S_subparietal_thickness | Cx Thickness | WIN | -0.002 | 0.002 | -1.071 | 619 | 0.284 | 0.564 | 0.002 |
| rh_S_temporal_inf_thickness | Cx Thickness | WIN | -0.002 | 0.002 | -1.226 | 619 | 0.221 | 0.492 | 0.002 |
| rh_S_temporal_sup_thickness | Cx Thickness | WIN | -0.002 | 0.002 | -1.512 | 619 | 0.131 | 0.367 | 0.004 |
| rh_S_temporal_transverse_thickness | Cx Thickness | WIN | -0.005 | 0.003 | -1.960 | 619 | 0.050 | 0.222 | 0.006 |
| rh_MeanThickness_thickness | Cx Thickness | WIN | -0.001 | 0.001 | -1.225 | 619 | 0.221 | 0.492 | 0.002 |
| lh_G.S_cingul.Ant_area | Cx Surface Area | WIN | -2.023 | 2.253 | -0.898 | 619 | 0.369 | 0.651 | 0.001 |
| lh_G.S_cingul.Mid.Post_area | Cx Surface Area | WIN | -0.093 | 1.259 | -0.074 | 618 | 0.941 | 0.970 | 0.000 |
| lh_G_cingul.Post.dorsal_area | Cx Surface Area | WIN | -0.274 | 0.792 | -0.345 | 619 | 0.730 | 0.889 | 0.000 |

| | | | | | | | | | |
|---|---|---|---|---|---|---|---|---|---|
| lh_G_cuneus_area | Cx Surface Area | WIN | 0.874 | 2.247 | 0.389 | 619 | 0.698 | 0.880 | 0.000 |
| lh_G_front_inf.Triangul_area | Cx Surface Area | WIN | 2.867 | 1.560 | 1.838 | 619 | 0.067 | 0.256 | 0.005 |
| lh_G_front_middle_area | Cx Surface Area | WIN | 2.532 | 4.800 | 0.527 | 619 | 0.598 | 0.838 | 0.000 |
| lh_G_Ins_lg.S_cent_ins_area | Cx Surface Area | WIN | -0.441 | 0.759 | -0.581 | 618 | 0.562 | 0.815 | 0.001 |
| ***lh_G_occipital_sup_area*** | ***Cx Surface Area*** | ***WIN*** | ***3.244*** | ***1.600*** | ***2.027*** | ***619*** | ***0.043*** | ***0.210*** | ***0.007*** |
| lh_G_orbital_area | Cx Surface Area | WIN | -1.620 | 1.907 | -0.849 | 619 | 0.396 | 0.678 | 0.001 |
| lh_G_pariet_inf.Angular_area | Cx Surface Area | WIN | -0.125 | 2.716 | -0.046 | 619 | 0.963 | 0.978 | 0.000 |
| lh_G_subcallosal_area | Cx Surface Area | WIN | 0.195 | 1.621 | 0.120 | 619 | 0.904 | 0.961 | 0.000 |
| lh_G_temp_sup.Plan_tempo_area | Cx Surface Area | WIN | 0.146 | 1.476 | 0.099 | 618 | 0.921 | 0.965 | 0.000 |
| lh_G_temporal_inf_area | Cx Surface Area | WIN | -1.029 | 2.981 | -0.345 | 619 | 0.730 | 0.889 | 0.000 |
| lh_Pole_occipital_area | Cx Surface Area | WIN | -0.295 | 1.768 | -0.167 | 619 | 0.868 | 0.951 | 0.000 |
| lh_S_central_area | Cx Surface Area | WIN | -0.152 | 2.488 | -0.061 | 618 | 0.951 | 0.971 | 0.000 |
| lh_S_collat_transv_ant_area | Cx Surface Area | WIN | -2.458 | 1.365 | -1.801 | 619 | 0.072 | 0.266 | 0.005 |
| lh_S_interm_prim.Jensen_area | Cx Surface Area | WIN | -0.477 | 1.352 | -0.353 | 619 | 0.724 | 0.889 | 0.000 |
| lh_S_oc.temp_lat_area | Cx Surface Area | WIN | -0.475 | 1.359 | -0.349 | 619 | 0.727 | 0.889 | 0.000 |
| lh_S_oc.temp_med.Lingual_area | Cx Surface Area | WIN | -1.603 | 1.838 | -0.872 | 618 | 0.384 | 0.664 | 0.001 |
| lh_S_orbital_lateral_area | Cx Surface Area | WIN | 1.029 | 0.729 | 1.412 | 619 | 0.158 | 0.406 | 0.003 |
| lh_S_orbital_med.olfact_area | Cx Surface Area | WIN | -0.850 | 0.654 | -1.300 | 619 | 0.194 | 0.458 | 0.003 |
| ***lh_S_orbital.H_Shaped_area*** | ***Cx Surface Area*** | ***WIN*** | ***-3.826*** | ***1.345*** | ***-2.845*** | ***619*** | ***0.005*** | ***0.091*** | ***0.013*** |
| ***lh_S_postcentral_area*** | ***Cx Surface Area*** | ***WIN*** | ***-6.998*** | ***3.415*** | ***-2.049*** | ***619*** | ***0.041*** | ***0.206*** | ***0.007*** |
| lh_S_subparietal_area | Cx Surface Area | WIN | -1.815 | 1.456 | -1.247 | 618 | 0.213 | 0.489 | 0.003 |
| lh_S_temporal_sup_area | Cx Surface Area | WIN | -4.025 | 5.218 | -0.771 | 619 | 0.441 | 0.716 | 0.001 |
| lh_WhiteSurfArea_area | Cx Surface Area | WIN | -80.132 | 73.520 | -1.090 | 619 | 0.276 | 0.561 | 0.002 |
| rh_G.S_occipital_inf_area | Cx Surface Area | WIN | -0.243 | 1.552 | -0.157 | 619 | 0.875 | 0.955 | 0.000 |
| rh_G_cuneus_area | Cx Surface Area | WIN | 0.168 | 2.265 | 0.074 | 619 | 0.941 | 0.970 | 0.000 |
| rh_G_front_inf.Triangul_area | Cx Surface Area | WIN | 2.358 | 1.497 | 1.575 | 618 | 0.116 | 0.347 | 0.004 |
| rh_G_occipital_sup_area | Cx Surface Area | WIN | 0.116 | 1.719 | 0.068 | 619 | 0.946 | 0.970 | 0.000 |
| ***rh_G_oc.temp_med.Parahip_area*** | ***Cx Surface Area*** | ***WIN*** | ***-3.692*** | ***1.733*** | ***-2.130*** | ***619*** | ***0.034*** | ***0.189*** | ***0.007*** |
| rh_G_orbital_area | Cx Surface Area | WIN | -2.074 | 2.026 | -1.024 | 619 | 0.306 | 0.587 | 0.002 |
| rh_G_precentral_area | Cx Surface Area | WIN | 2.416 | 2.237 | 1.080 | 619 | 0.281 | 0.562 | 0.002 |
| rh_G_temp_sup.Plan_tempo_area | Cx Surface Area | WIN | -0.590 | 0.995 | -0.593 | 619 | 0.553 | 0.810 | 0.001 |
| rh_Pole_occipital_area | Cx Surface Area | WIN | -1.297 | 3.056 | -0.424 | 619 | 0.671 | 0.871 | 0.000 |
| rh_S_central_area | Cx Surface Area | WIN | 0.623 | 2.440 | 0.255 | 619 | 0.799 | 0.924 | 0.000 |
| ***rh_S_circular_insula_inf_area*** | ***Cx Surface Area*** | ***WIN*** | ***-2.367*** | ***0.914*** | ***-2.591*** | ***619*** | ***0.010*** | ***0.111*** | ***0.011*** |
| rh_S_collat_transv_ant_area | Cx Surface Area | WIN | -2.036 | 1.682 | -1.211 | 619 | 0.226 | 0.494 | 0.002 |
| rh_S_collat_transv_post_area | Cx Surface Area | WIN | 1.199 | 0.825 | 1.454 | 619 | 0.147 | 0.387 | 0.003 |
| rh_S_occipital_ant_area | Cx Surface Area | WIN | -1.440 | 1.349 | -1.068 | 619 | 0.286 | 0.564 | 0.002 |
| rh_S_oc.temp_lat_area | Cx Surface Area | WIN | -2.625 | 1.699 | -1.545 | 619 | 0.123 | 0.357 | 0.004 |
| rh_S_oc.temp_med.Lingual_area | Cx Surface Area | WIN | -0.705 | 2.030 | -0.347 | 619 | 0.728 | 0.889 | 0.000 |
| ***rh_S_orbital.H_Shaped_area*** | ***Cx Surface Area*** | ***WIN*** | ***-3.894*** | ***1.358*** | ***-2.867*** | ***619*** | ***0.004*** | ***0.091*** | ***0.013*** |
| ***rh_S_postcentral_area*** | ***Cx Surface Area*** | ***WIN*** | ***-8.602*** | ***3.363*** | ***-2.558*** | ***619*** | ***0.011*** | ***0.111*** | ***0.010*** |
| ***rh_S_subparietal_area*** | ***Cx Surface Area*** | ***WIN*** | ***-3.704*** | ***1.761*** | ***-2.103*** | ***619*** | ***0.036*** | ***0.196*** | ***0.007*** |
| rh_S_temporal_inf_area | Cx Surface Area | WIN | -2.346 | 1.768 | -1.327 | 619 | 0.185 | 0.451 | 0.003 |
| lh_G.S_frontomargin_meancurv | Cx Curvature | WIN | 0.000 | 0.000 | -1.344 | 618 | 0.179 | 0.444 | 0.003 |
| lh_G.S_occipital_inf_meancurv | Cx Curvature | WIN | 0.000 | 0.000 | 0.203 | 619 | 0.839 | 0.939 | 0.000 |
| lh_G.S_paracentral_meancurv | Cx Curvature | WIN | 0.000 | 0.000 | -0.428 | 619 | 0.669 | 0.871 | 0.000 |
| lh_G.S_subcentral_meancurv | Cx Curvature | WIN | 0.000 | 0.000 | 1.450 | 619 | 0.148 | 0.387 | 0.003 |
| lh_G.S_cingul.Mid.Ant_meancurv | Cx Curvature | WIN | 0.000 | 0.000 | 1.141 | 619 | 0.254 | 0.534 | 0.002 |
| lh_G_cingul.Post.dorsal_meancurv | Cx Curvature | WIN | 0.000 | 0.000 | 1.905 | 618 | 0.057 | 0.244 | 0.006 |
| ***lh_G_cuneus_meancurv*** | ***Cx Curvature*** | ***WIN*** | ***0.000*** | ***0.000*** | ***2.161*** | ***618*** | ***0.031*** | ***0.180*** | ***0.008*** |
| lh_G_front_inf.Opercular_meancurv | Cx Curvature | WIN | 0.000 | 0.000 | 1.899 | 618 | 0.058 | 0.245 | 0.006 |
| lh_G_front_middle_meancurv | Cx Curvature | WIN | 0.000 | 0.000 | 0.488 | 618 | 0.625 | 0.843 | 0.000 |
| lh_G_front_sup_meancurv | Cx Curvature | WIN | 0.000 | 0.000 | 0.243 | 618 | 0.808 | 0.930 | 0.000 |
| lh_G_insular_short_meancurv | Cx Curvature | WIN | 0.000 | 0.000 | 0.742 | 619 | 0.458 | 0.730 | 0.001 |

| | | | | | | | | | |
|---|---|---|---|---|---|---|---|---|---|
| lh_G_occipital_middle_meancurv | Cx Curvature | WIN | 0.000 | 0.000 | 0.422 | 619 | 0.673 | 0.871 | 0.000 |
| lh_G_occipital_sup_meancurv | Cx Curvature | WIN | 0.000 | 0.000 | 0.463 | 619 | 0.643 | 0.847 | 0.000 |
| lh_G_oc.temp_lat.fusifor_meancurv | Cx Curvature | WIN | 0.000 | 0.000 | 1.232 | 619 | 0.218 | 0.492 | 0.002 |
| lh_G_oc.temp_med.Lingual_meancurv | Cx Curvature | WIN | 0.000 | 0.000 | 0.539 | 619 | 0.590 | 0.837 | 0.000 |
| lh_G_oc.temp_med.Parahip_meancurv | Cx Curvature | WIN | 0.000 | 0.000 | -1.006 | 619 | 0.315 | 0.595 | 0.002 |
| ***lh_G_orbital_meancurv*** | ***Cx Curvature*** | ***WIN*** | ***0.000*** | ***0.000*** | ***2.494*** | ***619*** | ***0.013*** | ***0.123*** | ***0.010*** |
| lh_G_pariet_inf.Supramar_meancurv | Cx Curvature | WIN | 0.000 | 0.000 | 1.477 | 619 | 0.140 | 0.377 | 0.004 |
| lh_G_parietal_sup_meancurv | Cx Curvature | WIN | 0.000 | 0.000 | -0.489 | 619 | 0.625 | 0.843 | 0.000 |
| lh_G_postcentral_meancurv | Cx Curvature | WIN | 0.000 | 0.000 | 0.084 | 619 | 0.933 | 0.969 | 0.000 |
| lh_G_precentral_meancurv | Cx Curvature | WIN | 0.000 | 0.000 | 0.504 | 617 | 0.615 | 0.843 | 0.000 |
| lh_G_rectus_meancurv | Cx Curvature | WIN | 0.000 | 0.000 | 1.218 | 619 | 0.224 | 0.493 | 0.002 |
| lh_G_subcallosal_meancurv | Cx Curvature | WIN | 0.000 | 0.000 | -0.848 | 619 | 0.397 | 0.678 | 0.001 |
| lh_G_temp_sup.G_T_transv_meancurv | Cx Curvature | WIN | 0.000 | 0.000 | 1.206 | 619 | 0.228 | 0.495 | 0.002 |
| lh_G_temp_sup.Plan_polar_meancurv | Cx Curvature | WIN | 0.000 | 0.000 | -0.722 | 619 | 0.471 | 0.733 | 0.001 |
| ***lh_G_temporal_inf_meancurv*** | ***Cx Curvature*** | ***WIN*** | ***0.000*** | ***0.000*** | ***2.516*** | ***619*** | ***0.012*** | ***0.123*** | ***0.010*** |
| lh_G_temporal_middle_meancurv | Cx Curvature | WIN | 0.000 | 0.000 | -1.247 | 618 | 0.213 | 0.489 | 0.003 |
| lh_Lat_Fis.ant.Horizont_meancurv | Cx Curvature | WIN | 0.000 | 0.000 | 1.728 | 619 | 0.084 | 0.289 | 0.005 |
| lh_Lat_Fis.ant.Vertical_meancurv | Cx Curvature | WIN | 0.000 | 0.000 | 0.459 | 617 | 0.647 | 0.848 | 0.000 |
| lh_Pole_temporal_meancurv | Cx Curvature | WIN | 0.000 | 0.000 | 1.564 | 617 | 0.118 | 0.353 | 0.004 |
| lh_S_central_meancurv | Cx Curvature | WIN | 0.000 | 0.000 | -0.509 | 618 | 0.611 | 0.843 | 0.000 |
| lh_S_cingul.Marginalis_meancurv | Cx Curvature | WIN | 0.000 | 0.000 | 1.547 | 619 | 0.122 | 0.357 | 0.004 |
| lh_S_circular_insula_ant_meancurv | Cx Curvature | WIN | 0.000 | 0.000 | 0.150 | 617 | 0.881 | 0.956 | 0.000 |
| lh_S_circular_insula_inf_meancurv | Cx Curvature | WIN | 0.000 | 0.000 | 0.469 | 619 | 0.639 | 0.847 | 0.000 |
| lh_S_circular_insula_sup_meancurv | Cx Curvature | WIN | 0.000 | 0.000 | -0.394 | 611 | 0.694 | 0.877 | 0.000 |
| lh_S_collat_transv_ant_meancurv | Cx Curvature | WIN | 0.000 | 0.000 | 1.201 | 618 | 0.230 | 0.496 | 0.002 |
| ***lh_S_front_inf_meancurv*** | ***Cx Curvature*** | ***WIN*** | ***0.000*** | ***0.000*** | ***2.594*** | ***618*** | ***0.010*** | ***0.111*** | ***0.011*** |
| lh_S_front_sup_meancurv | Cx Curvature | WIN | 0.000 | 0.000 | 0.536 | 618 | 0.592 | 0.837 | 0.000 |
| lh_S_interm_prim.Jensen_meancurv | Cx Curvature | WIN | 0.000 | 0.000 | -0.639 | 618 | 0.523 | 0.780 | 0.001 |
| lh_S_oc_middle.Lunatus_meancurv | Cx Curvature | WIN | 0.000 | 0.000 | 0.938 | 619 | 0.348 | 0.634 | 0.001 |
| lh_S_oc.temp_lat_meancurv | Cx Curvature | WIN | 0.000 | 0.000 | 0.172 | 619 | 0.863 | 0.948 | 0.000 |
| lh_S_oc.temp_med.Lingual_meancurv | Cx Curvature | WIN | 0.000 | 0.000 | 0.055 | 618 | 0.956 | 0.974 | 0.000 |
| lh_S_orbital_med.olfact_meancurv | Cx Curvature | WIN | 0.000 | 0.000 | 1.320 | 618 | 0.187 | 0.451 | 0.003 |
| lh_S_orbital.H_Shaped_meancurv | Cx Curvature | WIN | 0.000 | 0.000 | 1.321 | 618 | 0.187 | 0.451 | 0.003 |
| lh_S_precentral.inf.part_meancurv | Cx Curvature | WIN | 0.000 | 0.000 | 1.727 | 619 | 0.085 | 0.289 | 0.005 |
| lh_S_temporal_inf_meancurv | Cx Curvature | WIN | 0.000 | 0.000 | 1.135 | 618 | 0.257 | 0.534 | 0.002 |
| rh_G.S_paracentral_meancurv | Cx Curvature | WIN | 0.000 | 0.000 | -0.675 | 618 | 0.500 | 0.754 | 0.001 |
| rh_G.S_cingul.Ant_meancurv | Cx Curvature | WIN | 0.000 | 0.000 | -0.203 | 618 | 0.839 | 0.939 | 0.000 |
| rh_G.S_cingul.Mid.Ant_meancurv | Cx Curvature | WIN | 0.000 | 0.000 | 0.081 | 618 | 0.936 | 0.970 | 0.000 |
| rh_G.S_cingul.Mid.Post_meancurv | Cx Curvature | WIN | 0.000 | 0.000 | -0.204 | 619 | 0.838 | 0.939 | 0.000 |
| rh_G_cingul.Post.dorsal_meancurv | Cx Curvature | WIN | 0.000 | 0.000 | -0.316 | 617 | 0.752 | 0.898 | 0.000 |
| rh_G_cuneus_meancurv | Cx Curvature | WIN | 0.000 | 0.000 | 1.089 | 619 | 0.277 | 0.561 | 0.002 |
| rh_G_front_inf.Opercular_meancurv | Cx Curvature | WIN | 0.000 | 0.000 | -0.061 | 619 | 0.951 | 0.971 | 0.000 |
| ***rh_G_front_inf.Triangul_meancurv*** | ***Cx Curvature*** | ***WIN*** | ***0.000*** | ***0.000*** | ***2.259*** | ***619*** | ***0.024*** | ***0.163*** | ***0.008*** |
| rh_G_front_middle_meancurv | Cx Curvature | WIN | 0.000 | 0.000 | 1.152 | 618 | 0.250 | 0.527 | 0.002 |
| rh_G_front_sup_meancurv | Cx Curvature | WIN | 0.000 | 0.000 | 0.066 | 618 | 0.947 | 0.970 | 0.000 |
| rh_G_insular_short_meancurv | Cx Curvature | WIN | 0.000 | 0.000 | 1.343 | 618 | 0.180 | 0.444 | 0.003 |
| rh_G_occipital_sup_meancurv | Cx Curvature | WIN | 0.000 | 0.000 | -0.239 | 619 | 0.811 | 0.931 | 0.000 |
| ***rh_G_oc.temp_lat.fusifor_meancurv*** | ***Cx Curvature*** | ***WIN*** | ***0.000*** | ***0.000*** | ***2.176*** | ***619*** | ***0.030*** | ***0.180*** | ***0.008*** |
| rh_G_oc.temp_med.Parahip_meancurv | Cx Curvature | WIN | 0.000 | 0.000 | -1.672 | 619 | 0.095 | 0.311 | 0.004 |
| rh_G_orbital_meancurv | Cx Curvature | WIN | 0.000 | 0.000 | 0.634 | 619 | 0.526 | 0.783 | 0.001 |
| rh_G_pariet_inf.Angular_meancurv | Cx Curvature | WIN | 0.000 | 0.000 | -1.083 | 619 | 0.279 | 0.562 | 0.002 |
| rh_G_parietal_sup_meancurv | Cx Curvature | WIN | 0.000 | 0.000 | 0.301 | 619 | 0.764 | 0.905 | 0.000 |
| rh_G_precentral_meancurv | Cx Curvature | WIN | 0.000 | 0.000 | 0.726 | 618 | 0.468 | 0.733 | 0.001 |
| rh_G_rectus_meancurv | Cx Curvature | WIN | 0.000 | 0.000 | 1.299 | 618 | 0.194 | 0.458 | 0.003 |

| | | | | | | | | | |
|---|---|---|---|---|---|---|---|---|---|
| rh_G_subcallosal_meancurv | Cx Curvature | WIN | 0.000 | 0.000 | 0.119 | 618 | 0.905 | 0.961 | 0.000 |
| rh_G_temp_sup.G_T_transv_meancurv | Cx Curvature | WIN | 0.000 | 0.000 | 1.492 | 619 | 0.136 | 0.370 | 0.004 |
| rh_G_temporal_inf_meancurv | Cx Curvature | WIN | 0.000 | 0.000 | -0.090 | 618 | 0.928 | 0.968 | 0.000 |
| rh_G_temporal_middle_meancurv | Cx Curvature | WIN | 0.000 | 0.000 | 1.862 | 619 | 0.063 | 0.250 | 0.006 |
| rh_Lat_Fis.ant.Horizont_meancurv | Cx Curvature | WIN | 0.000 | 0.000 | 0.876 | 619 | 0.381 | 0.663 | 0.001 |
| ***rh_Pole_temporal_meancurv*** | ***Cx Curvature*** | ***WIN*** | ***0.000*** | ***0.000*** | ***2.428*** | ***619*** | ***0.015*** | ***0.134*** | ***0.009*** |
| rh_S_central_meancurv | Cx Curvature | WIN | 0.000 | 0.000 | -0.719 | 618 | 0.473 | 0.733 | 0.001 |
| ***rh_S_circular_insula_ant_meancurv*** | ***Cx Curvature*** | ***WIN*** | ***0.000*** | ***0.000*** | ***1.984*** | ***618*** | ***0.048*** | ***0.222*** | ***0.006*** |
| rh_S_circular_insula_inf_meancurv | Cx Curvature | WIN | 0.000 | 0.000 | 0.185 | 619 | 0.853 | 0.947 | 0.000 |
| rh_S_circular_insula_sup_meancurv | Cx Curvature | WIN | 0.000 | 0.000 | 1.623 | 615 | 0.105 | 0.328 | 0.004 |
| rh_S_collat_transv_ant_meancurv | Cx Curvature | WIN | 0.000 | 0.000 | 1.824 | 619 | 0.069 | 0.260 | 0.005 |
| ***rh_S_front_inf_meancurv*** | ***Cx Curvature*** | ***WIN*** | ***0.000*** | ***0.000*** | ***3.047*** | ***619*** | ***0.002*** | ***0.076*** | ***0.015*** |
| rh_S_interm_prim.Jensen_meancurv | Cx Curvature | WIN | 0.000 | 0.000 | 0.483 | 618 | 0.629 | 0.845 | 0.000 |
| rh_S_oc_middle.Lunatus_meancurv | Cx Curvature | WIN | 0.000 | 0.000 | 0.173 | 619 | 0.863 | 0.948 | 0.000 |
| rh_S_occipital_ant_meancurv | Cx Curvature | WIN | 0.000 | 0.000 | 1.303 | 619 | 0.193 | 0.458 | 0.003 |
| rh_S_oc.temp_lat_meancurv | Cx Curvature | WIN | 0.000 | 0.000 | -0.681 | 619 | 0.496 | 0.750 | 0.001 |
| rh_S_oc.temp_med.Lingual_meancurv | Cx Curvature | WIN | 0.000 | 0.000 | 1.592 | 618 | 0.112 | 0.343 | 0.004 |
| rh_S_orbital_lateral_meancurv | Cx Curvature | WIN | 0.000 | 0.000 | -0.183 | 619 | 0.855 | 0.947 | 0.000 |
| rh_S_orbital_med.olfact_meancurv | Cx Curvature | WIN | 0.000 | 0.000 | 0.351 | 616 | 0.726 | 0.889 | 0.000 |
| rh_S_orbital.H_Shaped_meancurv | Cx Curvature | WIN | 0.000 | 0.000 | 0.527 | 617 | 0.598 | 0.838 | 0.000 |
| rh_S_pericallosal_meancurv | Cx Curvature | WIN | 0.000 | 0.000 | -0.356 | 619 | 0.722 | 0.889 | 0.000 |
| rh_S_precentral.inf.part_meancurv | Cx Curvature | WIN | 0.000 | 0.000 | -0.751 | 618 | 0.453 | 0.725 | 0.001 |
| rh_S_precentral.sup.part_meancurv | Cx Curvature | WIN | 0.000 | 0.000 | 0.871 | 619 | 0.384 | 0.664 | 0.001 |
| rh_S_temporal_inf_meancurv | Cx Curvature | WIN | 0.000 | 0.000 | 1.498 | 619 | 0.135 | 0.368 | 0.004 |

**Supplementary Table 3. Win-by-Age-by-MoCA Interactions in metrics meeting p-fdr<0.05 for target effects (Tables 2&3)**

| Region, Measure | effect | beta | SE | t | df | p | paritalr2 |
|---|---|---|---|---|---|---|---|
| L Paraphippocampal White Matter, Volume | WIN*Age*MoCA | -0.0092 | 0.0042 | -2.1765 | 613 | 0.0299 | 0.0077 |
| L Inferior Lateral Ventricle, Volume | WIN*Age*MoCA | 0.0079 | 0.0043 | 1.8384 | 607 | 0.0665 | 0.0055 |
| R Inferior Lateral Ventricle, Volume | WIN*Age*MoCA | 0.0075 | 0.0043 | 1.7287 | 610 | 0.0844 | 0.0049 |
| L Middle Occipital Sulcus and Lunatus, Thickness | WIN*Age*MoCA | 0.0000 | 0.0000 | -1.5116 | 613 | 0.1312 | 0.0037 |
| L Heschl's gyrus, Thickness | WIN*Age*MoCA | 0.0000 | 0.0000 | -1.4874 | 614 | 0.1374 | 0.0036 |
| R Anterior Cingulate Gyrus and Sulcus, Thickness | WIN*Age*MoCA | 0.0000 | 0.0000 | -1.3811 | 614 | 0.1678 | 0.0031 |
| R Occpital Pole, Thickness | WIN*Age*MoCA | 0.0000 | 0.0000 | -1.3111 | 614 | 0.1903 | 0.0028 |
| L Superior Occipital Sulcus and Transversal, Thickness | WIN*Age*MoCA | 0.0000 | 0.0000 | -1.2793 | 613 | 0.2013 | 0.0027 |
| L Calcarine Sulcus, Thickness | WIN*Age*MoCA | 0.0000 | 0.0000 | -1.2249 | 614 | 0.2211 | 0.0024 |
| L Superior Occipital Gyrus, Thickness | WIN*Age*MoCA | 0.0000 | 0.0000 | -0.5036 | 614 | 0.6147 | 0.0004 |
| L Occpital Pole, Thickness | WIN*Age*MoCA | 0.0000 | 0.0000 | 0.4678 | 614 | 0.6401 | 0.0004 |
| R Fusiform White Matter, Volume | WIN*Age*MoCA | -0.0064 | 0.0157 | -0.4076 | 614 | 0.6837 | 0.0003 |
| L Mid Pos Cingulate Gyrus & Sulcus, Mean Curvature | WIN*Age*MoCA | 0.0000 | 0.0000 | -0.4035 | 614 | 0.6867 | 0.0003 |
| R Rectus Gyrus, Thickness | WIN*Age*MoCA | 0.0000 | 0.0000 | -0.3882 | 613 | 0.6980 | 0.0002 |
| L Middle Occipital Gyrus, Thickness | WIN*Age*MoCA | 0.0000 | 0.0000 | 0.2742 | 614 | 0.7841 | 0.0001 |
| L Cuneus Gyrus, Thickness | WIN*Age*MoCA | 0.0000 | 0.0000 | -0.1602 | 614 | 0.8728 | 0.0000 |
| L Entorhinal White Matter, Volume | WIN*Age*MoCA | -0.0002 | 0.0044 | -0.0419 | 614 | 0.9666 | 0.0000 |
| R Middle Temporal White Matter, Volume | WIN*Age*MoCA | -0.0007 | 0.0171 | -0.0404 | 614 | 0.9678 | 0.0000 |
